# Centrosome Migration and Apical Membrane Formation in Polarized Epithelial Cells: Insights from the MDCK Cyst Model

**DOI:** 10.1101/2024.06.17.598507

**Authors:** Po-Kai Wang, Keng-Hui Lin, Tang K. Tang

## Abstract

Polarization is crucial for the proper functioning of epithelial cells. Early hallmarks include the trafficking and enrichment of polarity molecules to form the apical membrane (AM) or cell -cell junctions, and the apical positioning of the centrosome. However, the dependencies among polarity molecules, AM formation, and centrosome positioning remain poorly understood. When cultured in Matrigel, *de novo* polarization of a single epithelial cell is often coupled with cytokinesis. During mitotic exit, centrosomes move to the future AM site, raising questions about their role in polarization. We perturbed centrosomes and polarity regulators in Matrigel-cultured cells and manipulated polarity direction using suspension culture to examine the relationships among polarization features. Surprisingly, centrosomal microtubules may not be necessary for centrosome positioning or AM formation, but their absence reduces the efficiency of AM formation. The polarity regulator Par3, rather than AM component trafficking, influences centrosome positioning. In suspension culture, centrosome migrate in the direction opposite to AM formation. Taken together, our findings define the hierarchical relationships among several polarization features and show that centrosome-based polarity is not universal in epithelial cells, providing new insights into the mechanisms of epithelial polarization.

**Impact statement:** Single-cell observations refute the expectation that centrosome positioning is a critical factor in the polarization of epithelial cells, while showing that proper centrosome positioning enhances the efficiency of cell polarization.

## Introduction

Cell polarity refers to spatial asymmetry in the shape, organization, and function of a cell. The development of cell polarity is a fundamental process in developmental and physiological biology, and aberrations in cell polarization can contribute to various developmental disorders and cancer (Hakanen et al., 2019; Halaoui & McCaffrey, 2015; Murdoch et al., 2003; Royer & Lu, 2011; Schneeberger et al., 2018; Wilson, 2011). Although polarizations in different cell types present in different morphologies, they share evolutionarily conserved machinery components that mediate these processes. For example, the partitioning defective (PAR) complexes, Crumbs complexes, and Scribble complexes are known to be asymmetrically occupied in different cellular compartments to regulate cell polarity (Goldstein & Macara, 2007; Iden & Collard, 2008; St Johnston & Ahringer, 2010). Furthermore, the centrosome is always located at a specific position, resulting in the centrosome–nucleus axis aligning with the polarization axis, which may be crucial for various types of polarized cells (Burakov & Nadezhdina, 2020; Tang & Marshall, 2012).

The centrosome is a distinct organelle in cells, composed of a centriole pair surrounded by a multi - protein cloud called pericentriolar material (PCM) (Conduit et al., 2015). The centrioles in the pair exhibit structural and functional asymmetry based on their generation, with the older, fully mature mother centriole carrying characteristic subdistal appendages (SDAs) and distal appendages (DAs) (Bornens, 2012; Gomes Pereira et al., 2021). The SDAs anchor microtubules to the mother centriole (Tateishi et al., 2013), while the DAs allow the mother centriole to dock to the plasma membrane (Tanos et al., 2013). The PCM comprises hundreds of proteins, including the scaffold protein pericentrin (PCNT) and the γ-tubulin ring complex, which are required for microtubule nucleation (Fong et al., 2008; Zimmerman et al., 2004). Because of its strong ability to nucleate microtubules, the centrosome functions as the primary microtubule organizing center and is considered to play a crucial role in regulating cell polarity (Bornens, 2012; de Anda et al., 2005). Laser ablation of the pericentrosomal region alters the polarized cell shape and affects cell migration (Wakida et al., 2010). Light inactivation or depletion of centrosomal proteins disrupts axon formation and neuronal migration (de Anda et al., 2010). Depletion of Par3 and dynein indirectly interferes with centrosome positioning and causes polarity defects in migrating cells (Schmoranzer et al., 2009). However, researchers have reported contradictory results in some cases. For instance, axons continue to grow when the centrosome in rodent hippocampal neurons is disrupted via laser ablation (Stiess et al., 2010). Moreover, the treatment of human umbilical vein endothelial cells (HUVECs) with centrinone, an inhibitor of centriole duplication (Wong et al., 2015), to deplete centrosomes does not affect the polarization of endothelial sprouting or cell migration. Notably, under conditions lacking non-centrosomal microtubules, centrosome depletion partially restored cell polarization, suggesting that under such conditions, the centrosome instead hinders cell polarization (Martin et al., 2018). Therefore, the exact role of the centrosome in cell polarity requires further clarification.

In polarized epithelial cells, the centrosome is localized at the apical region during interphase (Buckley & St Johnston, 2022; Ching et al., 2022; Rodriguez-Boulan & Macara, 2014), which contributes to the construction of an asymmetric microtubule network conducive to polarized vesicle trafficking (Feldman & Priess, 2012; Meiring et al., 2020). Establishing apical-basal polarity in epithelial cells requires the correct intracellular transportation of different polarity molecules to distinct domains of the plasma membrane (Jewett & Prekeris, 2018; Mellman & Nelson, 2008; Roman-Fernandez & Bryant, 2016). These processes have been extensively studied using *in vitro* three-dimensional (3D) cultures of vertebrate epithelial cells, such as Madin-Darby Canine Kidney (MDCK) cells seeded in the extracellular matrix (ECM), which spontaneously form polarized cysts with central lumens to mimic tubular structures *in vivo*. According to this model, previous studies have found that the orientation of apical-basal polarity depends on ECM-mediated signaling. In suspended culture without ECM or inhibition of ECM protein laminin, its receptor β1-integrin, or downstream signaling molecule small GTPase Rac1, polarity reversal occurs, leading to failure in lumen formation (O’Brien et al., 2001; Yu et al., 2005). Both integrin and Rac1 are necessary for laminin accumulation around the cyst and for inhibiting the RhoA-ROCK-mediated actomyosin on the basal side, which then enables the internalization of apical membrane proteins from the basolateral membrane (Bryant et al., 2014; Yu et al., 2008). Following ECM signaling, the well-known apical membrane marker Gp135 (also known as podocalyxin) begins to be internalized from the cell membrane into Rab11a-positive recycling endosomes when cells aggregate together or when a single cell undergoes cell division. These Gp135-positive vesicles containing Crumbs3 (Crb3) and Cdc42 are then delivered and tethered to the cell-cell contact site or cytokinesis site. Accumulation of these apical determinants forms the apical membrane initiation site (AMIS). Expansion of the apical membrane from the AMIS, combined with fluid pumping, generates a single lumen at the center of MDCK cysts (Bryant et al., 2010; Chou et al., 2016; Galvez-Santisteban et al., 2012; Klinkert et al., 2016; Mangan et al., 2016; Schluter et al., 2009).

Although numerous reports have been on the molecular networks involved in the *de novo* generation of the apical membrane and lumen, the sequential recruitment of apical determinants and the relative timing of centrosome positioning remain poorly characterized. The differences between cell-aggregation-derived polarization and cell-division-derived polarization are still critical in this field (Liang et al., 2022; Rathbun et al., 2020). Moreover, the role of apical migration of centrosomes in epithelial polarization remains unclear. Interestingly, previous studies have demonstrated that centrosomes can move toward the cytokinetic bridge during the late stages of cell division (Jonsdottir et al., 2010; Krishnan et al., 2022; Piel et al., 2001). Therefore, it is worthwhile to understand the coupling between the apical migration of centrosomes and cytokinesis.

In this work, we employed three major polarity indicators: (1) apical membrane protein recruitment, defined by Gp135 accumulation (Meder et al., 2005; Yu et al., 2007), (2) centrosome positioning, defined by the angles and relative distance between the nucleus and centrosome (Burute et al., 2017; Rodriguez-Fraticelli et al., 2012), and (3) organization of polarity regulators such as Par3, which controls tight junction assembly to partition the apical surface from the basolateral surface (Chen & Macara, 2005; Horikoshi et al., 2009). Here, we elucidate the detailed coupling between these polarity indicators during cytokinesis-induced *de novo* epithelial polarization in Matrigel and other cell culture conditions, which allow us to impose different polarity states. Our findings indicate that cytokinesis-induced centrosome positioning promotes epithelial polarization. The tight-junction-localized component Par3 is considered the upstream regulator of centrosome positioning and polarized vesicle trafficking. The trafficking of apical membrane components is downstream of centrosome positioning. The position of the centrosome is not sensitive to the disruption of individual centrosome components. Migrating centrosomes promote Gp135 accumulation during *de novo* epithelial polarization. Our study clarifies the relationship between different polarity indicators and the role of the centrosome in epithelial polarization.

## Results

### Apical membrane component proteins are recruited to the centrosome during *de novo* epithelial polarization

To capture the centrosome position and the distribution of Gp135 vesicles, we performed immunostaining at several stages in the transition from a single MDCK cell to a two-cell cyst. As a single non-polarized MDCK cell is embedded into Matrigel, most Gp135 proteins cluster around the interphase centrosome (Figure 1A, i) and strongly co-localize with the Rab11a-labeled recycling endosomes (Figure 1—figure supplement 1A). During mitosis, the Gp135 proteins are well dispersed in the cytosol at metaphase (Figure 1A, ii), weakly cluster near the centrosomes in early telophase (Telo, Figure 1A, iii), and gradually accumulate at the centrosomes from late telophase to the stage of cytokinetic bridge pre-abscission (pre-Abs) (Figure 1A, iii-iv), when the acetyl-tubulin-labeled cytokinetic bridge condenses significantly. Cytokinesis in most animal cells begins in the middle point of anaphase and ends shortly after the completion of mitosis in telophase. Cytokinetic abscission leads to the physical cut of the cytokinetic bridge connecting the daughter cells and concludes cell division. After cytokinetic bridge abscission (post-cytokinesis), the Gp135-positive vesicles are transported to the middle membrane region, which will form the AMIS between the two daughter cells (Figure 1A, v). Subsequently, exocytosis continues to expand the apical membrane and produces a lumen structure (Figure 1A, vi). The relative Gp135 signals near the centrosome regions of different stages show a clear maximum at the cytokinetic pre-abscission stage (Figure 1A–C). Notably, before bridge abscission, we detected a stronger signal of Gp135-positive vesicles surrounding the centrosomes compared to the cytosol and cytokinetic bridge (Figure 1A–B and D).

**Figure 1:**
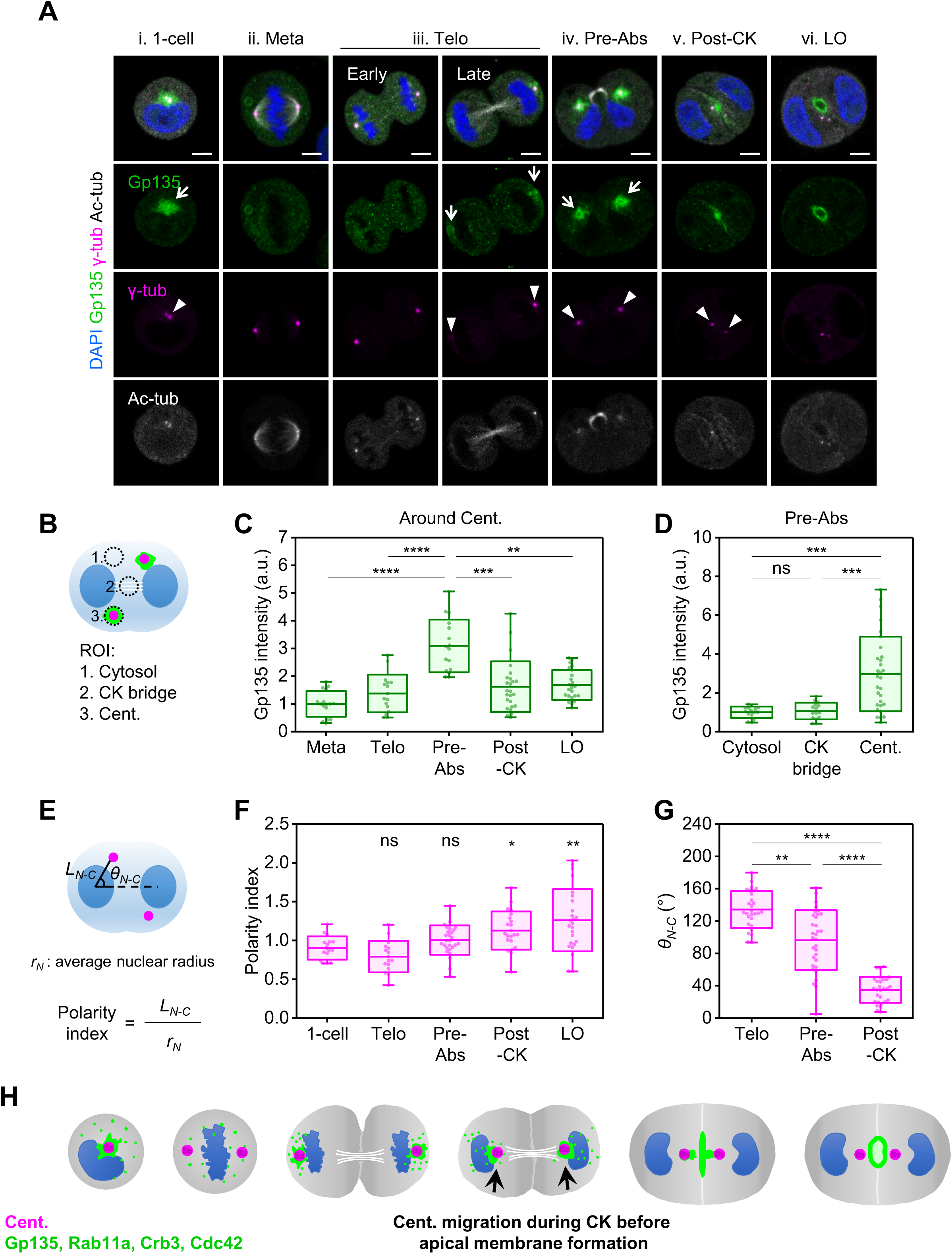
Localization of apical membrane protein Gp135 around the centrosome and its transport during epithelial polarization. (A) Single MDCK cells were cultured in Matrigel for 12 h. Immunostaining was performed with the following markers: apical membrane marker glycoprotein 135 (Gp135, green), centrosome marker γ-tubulin (magenta), acetyl-tubulin (white), and DAPI for nuclei (blue). Single confocal sections through the middle of a cyst are shown. The order of polarization is arranged from single cell (1-cell), metaphase (Meta), telophase (Telo), cytokinetic pre-abscission (Pre-Abs), post-cytokinesis (Post-CK), to lumen open (LO). Arrows indicate Gp135 clusters around centrosomes, while arrowheads point to centrosome positions. Scale bar: 5 μm. (B) Illustration showing the use of a fixed-size oval tool to select and measure Gp135 intensity in the chosen region. 1. Cytosol. 2. Cytokinesis (CK) bridge. 3. Centrosome (Cent.). (C–D) Boxplots of Gp135 intensity surrounding centrosomes at different stages normalized to the mean in metaphase (C) and at different cellular regions at the cytokinetic pre-abscission stage normalized to the mean in the cytosol (D). We analyzed >15 cells for each stage or region in three independent experiments. Statistical analyses were performed via one-way ANOVA and Dunn’s multiple comparisons (****p<0.0001, ***p<0.001, **p<0.01, ns: not significant). The midlines and boxes show the mean ± SD, with whiskers indicating minimum and maximum values (a.u., arbitrary units). (E) Illustration showing how the polarity index (*L_N-C_* [nucleus–centrosome distance] divided by *r_N_* [average nuclear radius]) and *θ_N-C_* (angle between the N-C and N-N axes) are calculated. (F–G) Boxplots depicting centrosome positions at different stages, shown by the polarity index (F) and *θ_N-C_* (G). We analyzed >16 cells per measurement in three independent experiments. Statistical analyses used one-way ANOVA and Dunn’s multiple comparisons (*p<0.05, **p<0.01, ****p<0.0001, ns: not significant). Midlines and boxes show the mean ± SD, with whiskers indicating minimum and maximum values. (H) Localization overview summarizing the localization of the centrosome (magenta) and recycling endosome/apical membrane components (green) during cell-division-directed polarization in Matrigel culture. White lines depict intercellular bridge microtubules, and nuclei are shown in blue. Arrows indicate apical membrane component localization around centrosomes and their transport from the centrosome to the AMIS.

We quantify the degree of polarization by the centrosome positions using the following quantities. The center point between two nuclei is set as the origin (O), representing the cyst center. The polarity index is defined as the nucleus (N)–centrosome (C) distance, *L_N-C_*, normalized by the average nuclear radius, *r_N_*. It quantifies the spatial displacement of the centrosome relative to the nucleus; the higher the value, the greater the degree of cell polarization. In addition, the nucleus–centrosome orientation (*θ_N-C_,* the angle between the N-C and N-N axes) characterizes the movement of the centrosome (Figure 1E, see Materials and methods) (Burute et al., 2017; Rodriguez-Fraticelli et al., 2012). Compared to the non-polarized 1-cell stage, the polarity index gradually increases from the cytokinetic pre-abscission to the post-cytokinesis stage, reaching the highest level during the lumen-open (LO) stage (Figure 1F). A high *θ_N-C_* value indicates that the centrosomes are located at the outside of two nuclei, which also corresponds to a larger *L*_O-C_ (origin–centrosome distance) and *L_IC_*(inter-centrosomal distance) (Figure 1—figure supplement 1B–E). From telophase to post-cytokinesis, centrosomes move from outside to the inside of two nuclei, and thus *θ_N-C_* decreases (Figure 1G).

Because recycling endosomes play a crucial role in trafficking apical membrane components such as Gp135, we examined other apical membrane components such as Crb3, Cdc42, and aPKC (Bryant et al., 2010; Klinkert et al., 2016; Schluter et al., 2009). We found that aPKC appears to diffuse in the cytoplasm during the M phase and concentrates at the AMIS only after cytokinesis (Figure 1—figure supplement 1F). This result may be due to a lower aPKC concentration, contributing to a weak immunostaining signal. Nevertheless, the signals of enhanced green fluorescent protein (EGFP)-Crb3 and EGFP-Cdc42 in MDCK cells exhibited similar patterns near the centrosome during cytokinetic pre-abscission and were transported to the AMIS from the centrosome in the post-cytokinesis stage, identical to Gp135 (Figure 1—figure supplement 2A–B).

Taken together, our data suggest that the centrosome serves as a hub for the recruitment of apical recycling endosomes. Therefore, the prior migration of the centrosome toward the center of cell doublets may facilitate the trafficking of apical recycling endosomes to the AMIS (Figure 1H).

### Centrosomes exhibit robust migration toward the center of two daughter cells after anaphase onset before Gp135 targeting the AMIS

To precisely define the hierarchy of centrosome migration, apical membrane formation, and apical-basal polarity establishment, we performed live cell imaging using MDCK cells that expressed fluorescently labeled centrosome protein, PACT-mKO1 (the PACT domain is a conserved centrosomal targeting motif of PCNT) (Gillingham & Munro, 2000), and apical membrane protein, EGFP-Gp135, cultured in Matrigel. The onset of anaphase is set as 0:00 (h: min). Consistent with our fixation data, live cell imaging showed that EGFP-Gp135-positive vesicles did not surround PACT-labeled centrosomes during metaphase. When cleavage furrow ingression occurred, EGFP-Gp135-positive vesicles were recruited to a region surrounding the centrosome (Figure 2A, 0:10). The movement of these EGFP-Gp135-positive vesicles followed the same path as centrosome migration (Figure 2A, 0:20–0:40), with the vesicles finally fusing to the middle membrane of the cell doublet (Figure 2A, 0:55). A small patch of Gp135 was also observed in a region unrelated to the centrosome (Figure 2A, 0:10-0:30). Subsequently, this Gp135 was either directly transported to AMIS or fused with centrosome-associated Gp135 and transported together. Notably, this protein patch was only observed when Gp135 was overexpressed in the cells. When endogenous Gp135 proteins were stained (Figure 1A), this obvious protein patch was not observed, suggesting that overexpression of Gp135 protein may lead to an increased local concentration of it in this region.

**Figure 2:**
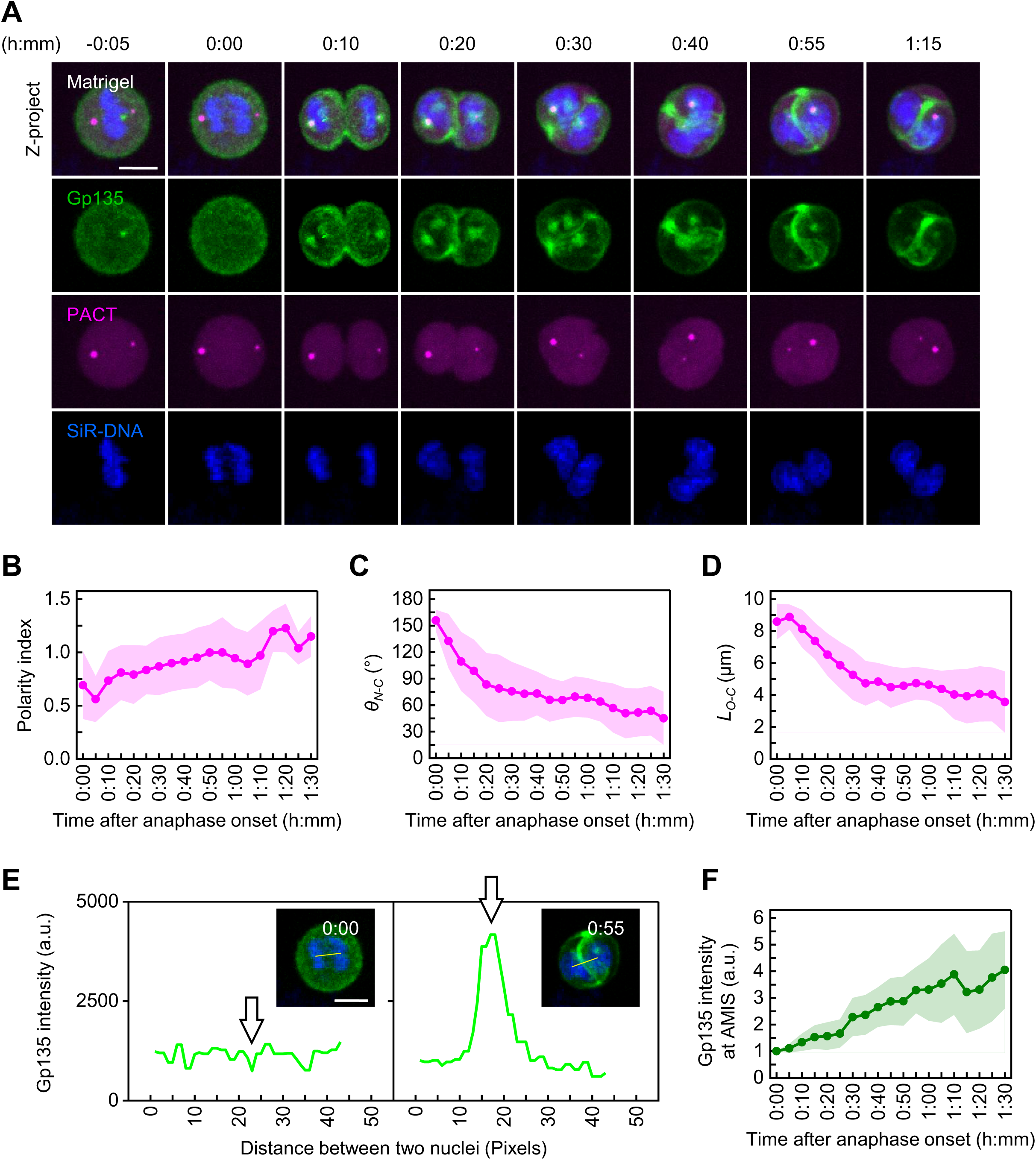
Timing of centrosome migration and Gp135 targeting to the AMIS. (A) Time-lapse snapshots of MDCK cells expressing EGFP-Gp135 (green) and PACT-mKO1 (magenta, centrosome marker) in Matrigel. Nuclei were labeled with SiR-DNA (blue) before live imaging. Z-projection images of a dividing cell are shown. Time stamps show hours and minutes, with 0:00 set at the first frame of anaphase onset. Scale bar: 10 μm. (B–D) Change in polarity index, *θ_N-C_*, and *L_O-C_*over time. Each data point represents the average at a given time (>10 cells in Matrigel culture from three independent experiments). The lines show the means, and the shaded regions indicate SD values. (E) Fluorescent profiles of EGFP-Gp135 along the line connecting the two nuclei in a cell doublet. White arrows indicate the central value used to indicate the level of Gp135 on the AMIS (a.u., arbitrary units). Time stamps show hours and minutes, with 0:00 set to the first frame after anaphase onset. Scale bar: 10 μm. (F) Change in the central value in the EGFP-Gp135 fluorescent profile over time. The value of each time point was normalized to the value at 0:00. Each data point represents the average fluorescent intensity at a given time (10 cells in Matrigel culture from three independent experiments). The line shows the mean, and the shaded region indicates SD values (a.u., arbitrary units).

Using our live cell movies, we measured the polarity index and found that cell polarization was approximately begun after the onset of anaphase (Figure 2B). *θ_N-C_* and *L*_O-C_ decreased significantly within 30 min after the anaphase onset and then plateaued (Figure 2C, D). These results suggest that centrosomes migrate toward the center of the cell doublet during the initial 30 min following the onset of anaphase, followed by a stable phase with reduced movement. Intriguingly, we observed this type of centrosome movement only in 3D culture cells embedded in Matrigel. In contrast, in adherent two-dimensional (2D) culture cells grown on a cover glass, the centrosomes exhibited random migration patterns (Figure 2—figure supplement 1A–C).

To investigate the temporal process of apical membrane formation, we measured the intensity profile of EGFP-Gp135 along the line connecting the two nuclei; we took the central value as an indicator of the amount of Gp135 on the AMIS (Figure 2E, white arrows). The intensity continually increased after the onset of anaphase (Figure 2F). As shown in Figure 2A, the EGFP-Gp135 signal at the middle membrane of the cell doublet increased significantly after 30 min from the onset of anaphase, coinciding with a reduced movement of the centrosomes (Figure 2C, D). These findings are consistent with our immunostaining results, indicating that centrosome migration precedes the targeting of Gp135 to the AMIS.

### Centrosomes influence the efficiency of early polarization but are not essential for late-stage lumen formation

To investigate the role of centrosomes in guiding Gp135-labeled vesicle trafficking for polarization, we depleted centrosomes in MDCK cells by applying centrinone, a reversible inhibitor of PLK4 kinase for centriole duplication in vertebrate cells. Centrinone treatment effectively blocked centriole duplication in MDCK cells and progressively depleted centriole numbers during cell cycle progression (Figure 3—figure supplement 1A, B). Three days after centrinone treatment, nearly 80% of cells lost their centrioles (Figure 3—figure supplement 1A), indicating the effective removal of centrosomes by centrinone treatment in MDCK cells.

**Figure 3:**
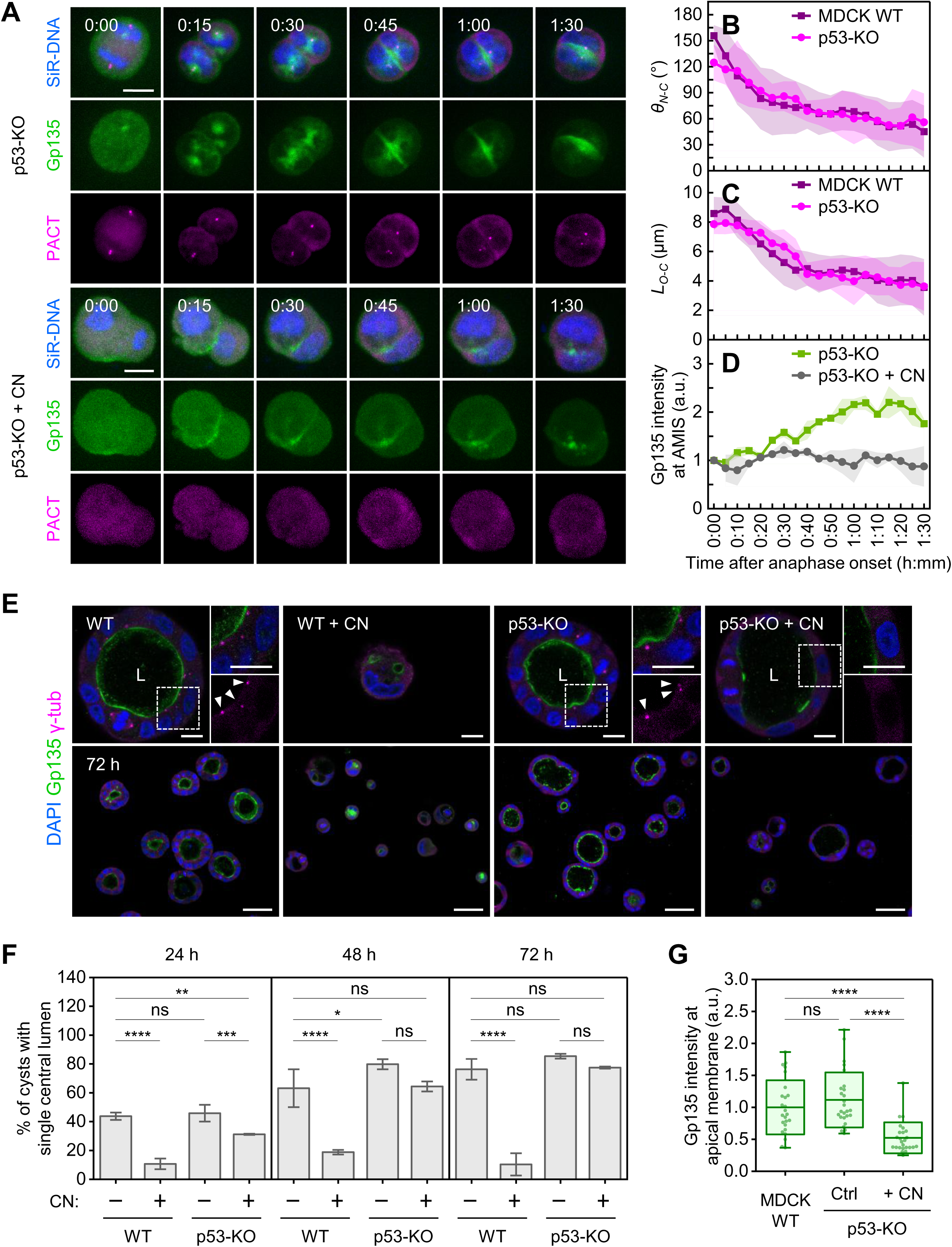
Impact of centriole depletion on apical lumen formation and Gp135 protein levels. (A) Time-lapse snapshots of p53-KO and centrinone (CN)-treated cells expressing EGFP-Gp135 (green) and PACT-mKO1 (magenta, centrosome marker) in Matrigel. Nuclei were labeled with SiR-DNA (blue) before live imaging. Z-projection images of dividing cells are shown. Time stamps show hours and minutes, with 0:00 set at the first frame of anaphase onset. Scale bar: 10 μm. (B–C) Change in *θ_N-C_* and *L_O-C_* over time. Each data point represents the average at a given time (>10 MDCK WT and >3 p53-KO cells in Matrigel culture from three independent experiments). The lines show the means, and the shaded regions indicate SD values. (D) Change in the central value of the EGFP-Gp135 fluorescent profile over time. The value of each time point was normalized to the value at 0:00. Each data point represents the average fluorescent intensity at a given time (n = 3 [p53-KO; olive] and 3 [p53-KO + CN; gray]). The lines show the means, and the shaded regions indicate SD values (a.u., arbitrary units). (E) Single MDCK WT and p53-KO cells, with or without CN treatment, after 72 h of Matrigel culture. Single confocal sections through the middle of cysts are shown with immunofluorescent signals of indicated markers: apical membrane Gp135 (green), centrosome marker γ-tubulin (magenta), and DAPI for nuclei (blue). "L" denotes the lumen. WT cells treated with CN remain at the single-cell stage. The insets display enlarged images of the region in the yellow box. Arrowheads indicate the presence of centrosomes. Scale bar: 10 μm. The bottom panels represent a larger view of the 3D culture. Scale bar: 50 μm. (F) Quantification of the proportion of MDCK cysts with a single central lumen after being cultured for different durations (24, 48, 72 h). The lumen structure was identified by Gp135 staining. n = 92 (24 h), 231 (48 h), 78 (72 h) cysts (MDCK WT); n = 22 (24 h), 133 (48 h), 59 (72 h) cysts (WT + CN); n = 154 (24 h), 170 (48 h), 204 (72 h) cysts (p53-KO); n = 105 (24 h), 128 (48 h), 149 (72 h) cysts (p53-KO + CN) analyzed for each measurement in three independent experiments. Statistical analyses used two-way ANOVA and Tukey multiple comparisons (*p<0.05, **p<0.01, ***p<0.001, ****p<0.0001, ns: not significant). Values represent the mean ± SD. (G) Boxplot of Gp135 intensity on the apical membrane normalized to the mean in WT cells. n = 25 (MDCK WT), 26 (p53-KO), 25 (p53-KO + CN) cysts were analyzed in three independent experiments. Statistical analyses were performed via one-way ANOVA and Dunn’s multiple comparisons (ns: not significant, ****p<0.0001). The midlines and boxes show the mean ± SD, with whiskers indicating minimum and maximum values (a.u., arbitrary units).

Previous studies have reported that a loss of centrosomes in cells activates p53, followed by activation of p21, which causes cell cycle arrest in non-transformed cells (Fong et al., 2016; Lambrus et al., 2016; Meitinger et al., 2016). However, cell division is required in our experimental setup to initiate epithelial polarization. To overcome the problem of cell cycle arrest, we generated a p53 knockout (KO) line of MDCK cells using the clustered regularly interspaced short palindromic repeats (CRISPR)/Cas9 editing system (Figure 3—figure supplement 1C, D). We found that the MDCK cells carry normal p53 activity and exhibit increased p21 protein levels after centrinone treatment (Figure 3—figure supplement 1E, F). As expected, the p53-KO MDCK cells do not induce p21 and continue proliferating after centrosome depletion (Figure 3—figure supplement 1F). Compared with wild-type (WT) cells, the p53-KO cells show no difference in polarity index, *θ_N-C_*, or *L*_O-C_ during *de novo* polarization (Figure 3A–C and Figure 3—figure supplement 1G), indicating that the loss of p53 does not interfere with centrosome movement.

Next, we treated the p53-KO MDCK cells with centrinone for three days, seeded them into Matrigel, and observed them by live cell imaging (Figure 3—figure supplement 1H). Our results showed that, in the absence of centrosomes, EGFP-Gp135 did not cluster in the cytoplasm at any time points (Figure 3A), and the accumulation of EGFP-Gp135 at the middle membrane was significantly reduced (Figure 3A, D). Together, our time-lapse imaging and centrosome depletion studies suggest that centrosome depletion causes a reduction in Gp135 accumulation at the AMIS due to a lack of Gp135 near the centrosome during cytokinesis. We then performed immunostaining, which supported our live cell imaging observations (Figure 3—figure supplement 2A, B).

We extended the observation time to 48 and 72 hours and found that more than 60% of centrosome-depleted p53-KO cells, which overcome the proliferation arrest, form cysts with a central lumen (Figure 3E, F). For both WT and p53-KO cells, the percentage of single-lumen cysts is higher than centrosome-depleted p53-KO cells at 24 hours, but the differences decrease over time (Figure 3E, F). However, the cell heights of the centrosome-depleted cysts are less uniform (Figure 3E, p53-KO + centrinone), and the Gp135 levels at the apical region remained lower than those of the controls (WT and p53-KO) (Figure 3G). Although p53-KO can overcome cell cycle arrest after centrosome depletion, its cell proliferation rate is still reduced (Figure 3—figure supplement 2C). We, therefore, compared the percentage of lumen formation in two-cell stage cysts to eliminate the effect of proliferation variation. 24 hours after seeding in Matrigel, more than 50% of WT and p53-KO two-cell cysts formed a single central lumen. In contrast, only 30% of centrinone-treated p53-KO two-cell cysts exhibited a single central lumen (Figure 3—figure supplement 2D). Taken together, our data suggest that centrosome depletion primarily affects the early steps of epithelial polarization but not the later steps of lumen formation.

### Disrupting centriole/centrosome structures and centrosomal microtubules does not alter centrosome positioning, but Gp135 trafficking is facilitated by centrosomal microtubules

Centrosomes are a large protein complex; thus, we sought to determine which components affect the position of centrosomes and the trafficking of Gp135 vesicles by perturbing centrosome components without completely removing the centrosome. We perturbed molecules associated with centriole/centrosome structures, such as CEP164 (an outer-layer protein of the DA), ODF2 (the base component of the SDA), and PCNT (the scaffold protein of the PCM) (Figure 4A). Each of these molecules has been reported to affect centrosome migration in different experimental systems (Hannaford et al., 2022; Hung et al., 2016; Krishnan et al., 2022; Pitaval et al., 2017).

**Figure 4:**
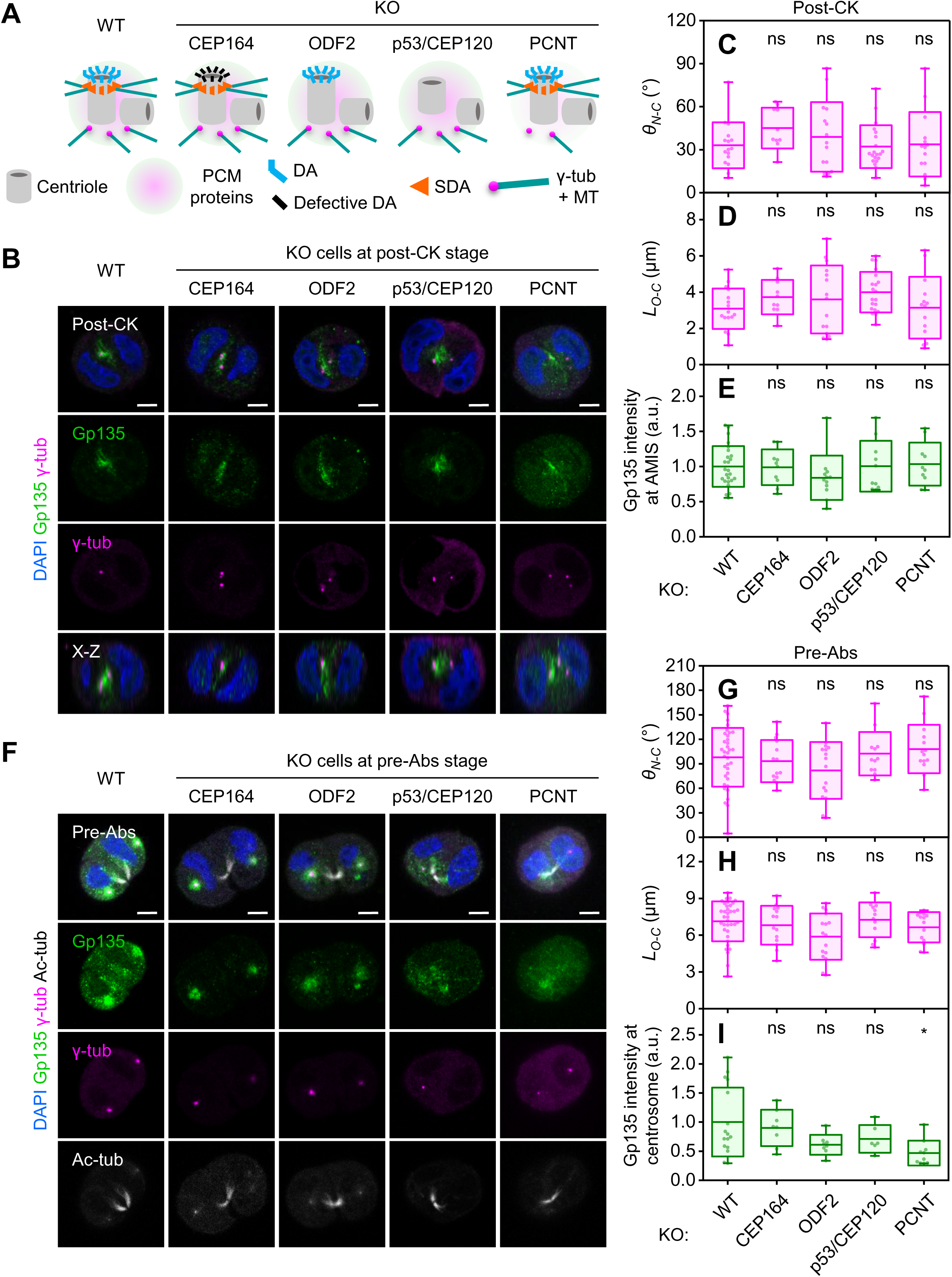
Centrosome migration in CEP164, ODF2, CEP120, and PCNT KO cells during epithelial polarization. (A) Illustration depicting individual centrosomal structures (DA, SDA, and PCM proteins) and the state of γ-tubulin and microtubules (γ-tub + MT) on the centrosome in cells with knockout (KO) of *CEP164*, *ODF2*, *p53/CEP120*, or *PCNT* genes. (B, F) Single MDCK cells of different genotypes were cultured in Matrigel for 12 h. Images show post-cytokinesis (CK) cells (B) or cells during pre-abscission (pre-Abs, F) with labeled markers: Gp135 (green), γ-tubulin (magenta), acetyl-tubulin (white), and DAPI for nuclei (blue). Images shown are single confocal sections through the middle of cells and side-view x-z cross-sections. Scale bar: 5 μm. (C–D, G–H) Boxplots of centrosome positions, represented by *θ_N-C_* and *L_O-C_*, for post-CK cells (C–D, n = 16 [WT], 10 [CEP164-KO], 15 [ODF2-KO], 20 [CEP120-KO], 14 [PCNT-KO]) and pre-Abs cells (G–H, n = 36 [WT], 14 [CEP164-KO], 16 [ODF2-KO], 12 [CEP120-KO], 14 [PCNT-KO]). Cells were analyzed in three independent experiments. Statistical analyses used one-way ANOVA and Dunn’s multiple comparisons (ns: not significant). Midlines and boxes show the mean ± SD, with whiskers indicating minimum and maximum values. (E, I) Boxplots of Gp135 intensity at the AMIS and centrosomes normalized to the mean in WT cells. Post-CK cell doublets: n = 25 (WT), 10 (CEP164-KO), 12 (ODF2-KO), 10 (CEP120-KO), and 10 (PCNT-KO). Pre-Abs cells: n = 16 (WT), 8 (CEP164-KO), 8 (ODF2-KO), 6 (CEP120-KO), and 10 (PCNT-KO). Data from three independent experiments. Statistical analyses were performed via one-way ANOVA and Dunn’s multiple comparisons (ns: not significant, *p<0.05). The midlines and boxes show the mean ± SD, with whiskers indicating minimum and maximum values (a.u., arbitrary units).

We generated KO lines of MDCK cells for CEP164, ODF2, and PCNT (Figure 4—figure supplement 1 and 2), as well as a KO line of CEP120 in the *p53^-/-^* background, since we were unable to generate the CEP120 KO line in p53-WT cells for unknown reasons (Figure 4—figure supplement 2E–H). CEP120 is an important regulator of centriole elongation (Comartin et al., 2013; Lin et al., 2013), and its KO enables the production of centrioles lacking both DA and SDA simultaneously (Figure 4A) (Tsai et al., 2019). Our results showed no significant differences in *θ_N-C_*or *L*_O-C_ between the WT and KO cells in the post-cytokinesis stage despite the presence of structural defects on the centriole/centrosome (Figure 4B–D). Gp135-labeled vesicles are still targeted to the center of cell doublets in the Matrigel culture (Figure 4E). Depletion of these centriole/centrosome structures does not affect polarized vesicle trafficking or centrosome positioning.

To determine whether defective centrosomes affect the timing of centrosome migration, we examined the centrosome position at the cytokinetic pre-abscission stage, as indicated by *θ_N-C_* and *L*_O-C_ (Figure 4F–H); at this stage, centrosomes have initiated migration but have not yet reached their final position (Figure 1A, G). We found defective centrosomes also initiated migration at the cytokinetic pre-abscission stage in various KO lines (Figure 4F). The *θ_N-C_* and *L*_O-C_ values showed no significant differences between the KO and WT cells (Figure 4G, H). These findings suggest that the timing of centrosome migration is not delayed in KO cells following the individual disruption of SDA, DA, or PCM components. However, during the pre-abscission stage, the intensity of Gp135 surrounding the centrosome is reduced in the PCNT-KO cells compared with the WT and other KO lines (Figure 4I), suggesting that centrosomal microtubules from PCM may play a role in recruiting Gp135 to the centrosome.

Microtubules serve as cables that generate mechanical forces facilitating centrosome migration (Laan et al., 2012; Okumura et al., 2018). These highlight the critical importance of the microtubule–centrosome connection. Furthermore, SDA has been implicated in anchoring microtubules to the centrosome (Tateishi et al., 2013). However, disruption of SDA through ODF2 knockout did not significantly affect centrosome positioning (Figure 4B–D, F–H), suggesting that SDA is not the sole determinant for centrosome migration. Additionally, although PCNT knockout cells show reduced microtubule nucleation ability (Figure 4—figure supplement 3A, B; Gavilan et al., 2018), they still recruit a small amount of γ-tubulin (Figure 4F). This observation suggests that the connection between microtubules and centrosome is not completely abolished.

To further investigate the role of centrosomal microtubules in centrosome positioning and polarized trafficking, we used a known method to disrupt centrosomal microtubule nucleation from the centrosome by expressing dominant-negative fragments of the γ-tubulin activator CDK5RAP2 and its binding protein NEDD1 (Vinopal et al., 2023). It was reported that exogenous expression of two DNA constructs encoding the centrosome-targeting carboxy-terminal domain (C-CTD) of CDK5RAP2 (Figure 4—figure supplement 3C) and the γ-tubulin-binding domain (gTBD) of NEDD1 (N-gTBD) (Figure 4—figure supplement 3D) in cells resulted in almost complete depletion of γ-tubulin and a marked reduction in centrosomal microtubule nucleation at the centrosome. Here, we co-expressed N-gTBD-mCherry and C-CTD-mCherry in MDCK cells to selectively inhibit centrosomal microtubules. Cells were then embedded in Matrigel and cultured for 24 hours to assess centrosome positioning. As expected, γ-tubulin dispersed from the centrioles, whereas acetylated tubulin and CEP120 (marking the centriole) could still be detected as distinct dots (Figure 4—figure supplement 3E, F, yellow arrowheads). Interestingly, the centriole dots were still remained positioned on the inner side of the two nuclei in cell doublets. Both the *θ_N-C_* and *L_O-C_* values showed no significant differences compared to those in control cells (Figure 4—figure supplement 3G, H), indicating that centrosomal microtubules may not be required for centrosome positioning. However, the intensity of Gp135 at the apical membrane were reduced at the two-cell stage (Figure 4—figure supplement 3I, J), suggesting that although centrosomal microtubules may not influence centrosome positioning, they enhance the efficiency of polarized trafficking and contribute to epithelial polarization.

### The polarity regulator Par3 first emerges at the cytokinesis site to regulate centrosome positioning and polarized vesicle trafficking during *de novo* epithelial polarization

We next examined molecules that have been previously reported to be involved in MDCK polarization (Bryant et al., 2010; Horikoshi et al., 2009; Martin-Belmonte et al., 2007; Roland et al., 2011; Schluter et al., 2009) to assess whether these molecules affect centrosome positioning. We first applied short hairpin RNA (shRNA) interference to deplete the molecules involved in vesicle trafficking, such as Rab11a, Sec15a, and myosin Vb (MyoVb) in WT cells (Figure 5—figure supplement 1A–C). Although depletion of the regulator of vesicle trafficking leads to a dispersion of Gp135, the centrosomes showed normal localization (Figure 5A–D). These results agree with our findings above, indicating that centrosome positioning governs the efficiency of polarized trafficking (Figure 3). Inhibition of vesicle transport mediated by Rab11a, Sec15a, or MyoVb does not affect centrosome positioning.

**Figure 5:**
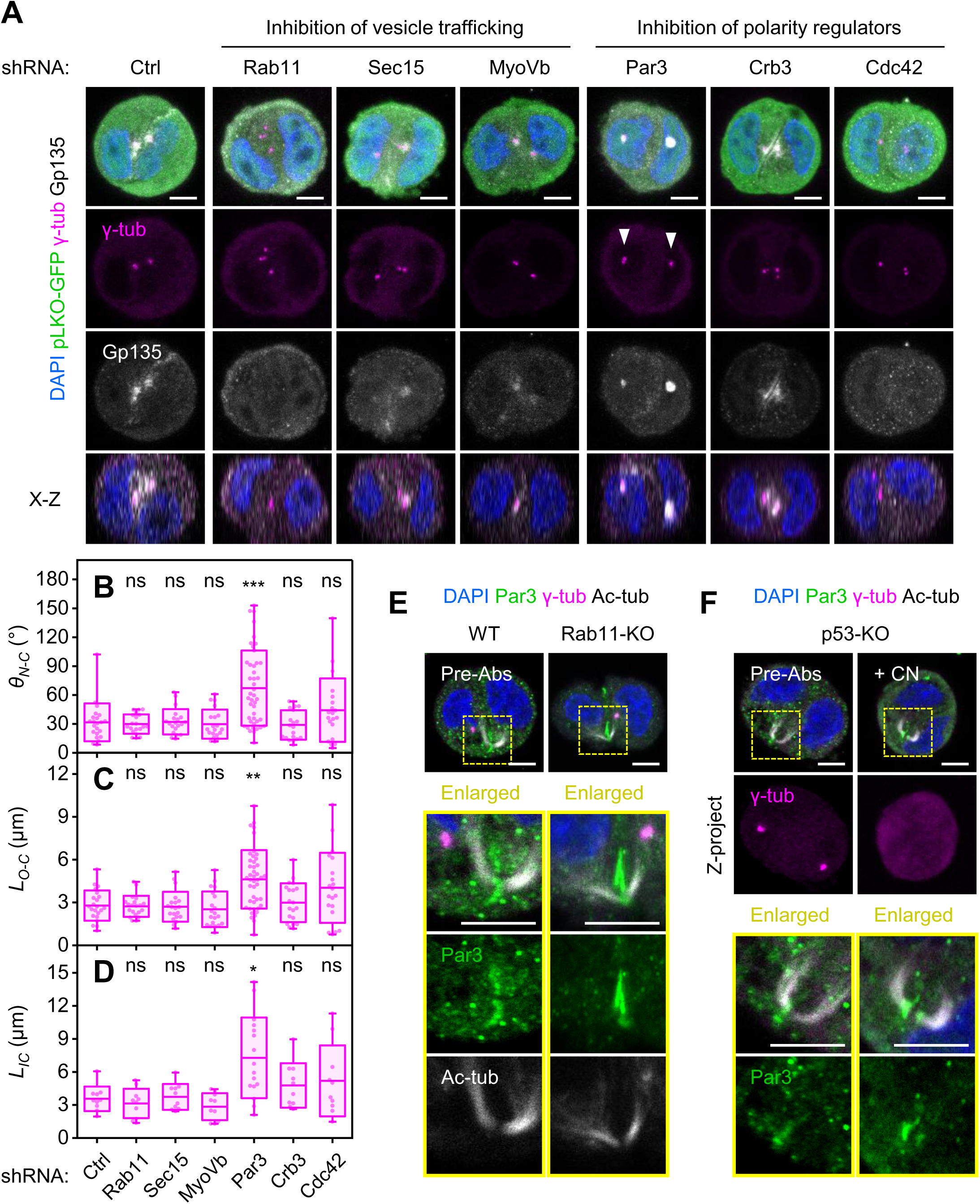
Par3 as the upstream regulator of centrosome migration and polarized vesicle transport. (A) Single MDCK cells, stably expressing pLKO-GFP-shRNA to knock down specified genes, cultured in Matrigel for 12 h. Images show post-cytokinesis cells with labeled markers: γ-tubulin (magenta), Gp135 (white), pLKO-GFP (green), and DAPI for nuclei (blue). Z-projection images between two centrosomes and the side-view x-z cross-section (bottom) are shown. "Ctrl" indicates the expression of scrambled-sequence shRNA. Arrowheads point to mislocalized centrosomes. Scale bar: 5 μm. (B–D) Boxplots of centrosome positions, represented by *θ_N-C_*, *L_O-C_*(B–C, n = 22 [sh-Ctrl], 20 [sh-Rab11a], 20 [sh-Sec15a], 20 [sh-MyoVb], 44 [sh-Par3], 20 [sh-Crb3], 20 [sh-Cdc42]), and *L_IC_* (D, n = 10 [sh-Ctrl], 10 [sh-Rab11a], 10 [sh-Sec15a], 10 [sh-MyoVb], 16 [sh-Par3], 10 [sh-Crb3], 10 [sh-Cdc42]), in post-cytokinesis cells. Cells were analyzed in three independent experiments. Statistical analyses used one-way ANOVA and Dunn’s multiple comparisons (ns: not significant, ***p<0.001, **p<0.01, *p<0.05). Midlines and boxes show the mean ± SD, with whiskers indicating minimum and maximum values. (E) Single WT and Rab11a-KO MDCK cells were cultured in Matrigel for 12 h with labeled markers: Par3 (green), γ-tubulin (magenta), acetyl-tubulin (white), and DAPI for nuclei (blue). Single confocal sections of pre-abscission cells are shown. The bridge region is enlarged from the yellow box. Scale bar: 5 μm. (F) Single confocal sections of p53-KO cells, with or without centrinone (CN) treatment, cultured in Matrigel for 12 h with labeled markers: Par3 (green), γ-tubulin (magenta), acetyl-tubulin (white), and DAPI for nuclei (blue). The bridge region shown in the yellow box is enlarged. The bottom panels display Z-projected images to demonstrate the complete depletion of centrosomes. Scale bar: 5 μm.

Next, we applied shRNA to deplete polarity regulators, such as Par3, Cdc42, and Crb3, in WT cells (Figure 5—figure supplement 1D–F). We found that Par3 depletion, but not Cdc42 or Crb3, significantly affects the centrosome positioning, as shown by a large deviation in *θ_N-C_*, *L_O-C_*, and *L_IC_* compared with the control (Figure 5A–D). The centrosomes do not move toward the AMIS after cytokinesis. Interestingly, we sometimes observed that in Par3-depleted cysts, Gp135- and Rab11a-positive vesicles are concentrated near mislocalized centrosomes in the post-cytokinesis stage (Figure 5A and Figure 5—figure supplement 1G). These data suggest that Par3 acts upstream of both centrosome positioning and polarized vesicle trafficking.

We next examined Par3 localization during *de novo* epithelial polarization through cytokinesis (Figure 5—figure supplement 1H). The antibody used can recognize PARD3 but not PARD3B, which are encoded by different genes located on chromosomes 2 and 37, respectively. In MDCK cells, this antibody signal was predominantly concentrated at tight junctions, with a very weak signal at the apical membrane. In Matrigel culture, our results show that Par3 first emerges at the cell-cell interface of the mitotic cleavage site at the cytokinetic pre-abscission stage (Figure 5—figure supplement 1H), in contrast to the apical membrane components, which concentrate around the centrosome at this stage (Figure 1A, B and Figure 1—figure supplement 2A, B). Indeed, we found that the cell-cell junction components Par3, actin, ZO-1, and E-cadherin did not surround migrating centrosomes; rather, they localized to the cytokinesis site and cell-cell contact interface at the pre-abscission stage (Figure 5—figure supplement 2A, B). The cell-cell junction components, especially Par3, are recruited to the center of the cell doublets before positioning the centrosomes and targeting apical membrane components to the AMIS. The spatial-temporal localization of proteins involved in epithelial polarization is summarized in Table 1. The sequential order of these events and our knockdown experiments (Figure 5A-D) suggest that Par3 acts upstream of centrosome positioning and polarized vesicle trafficking.

**Table 1.**
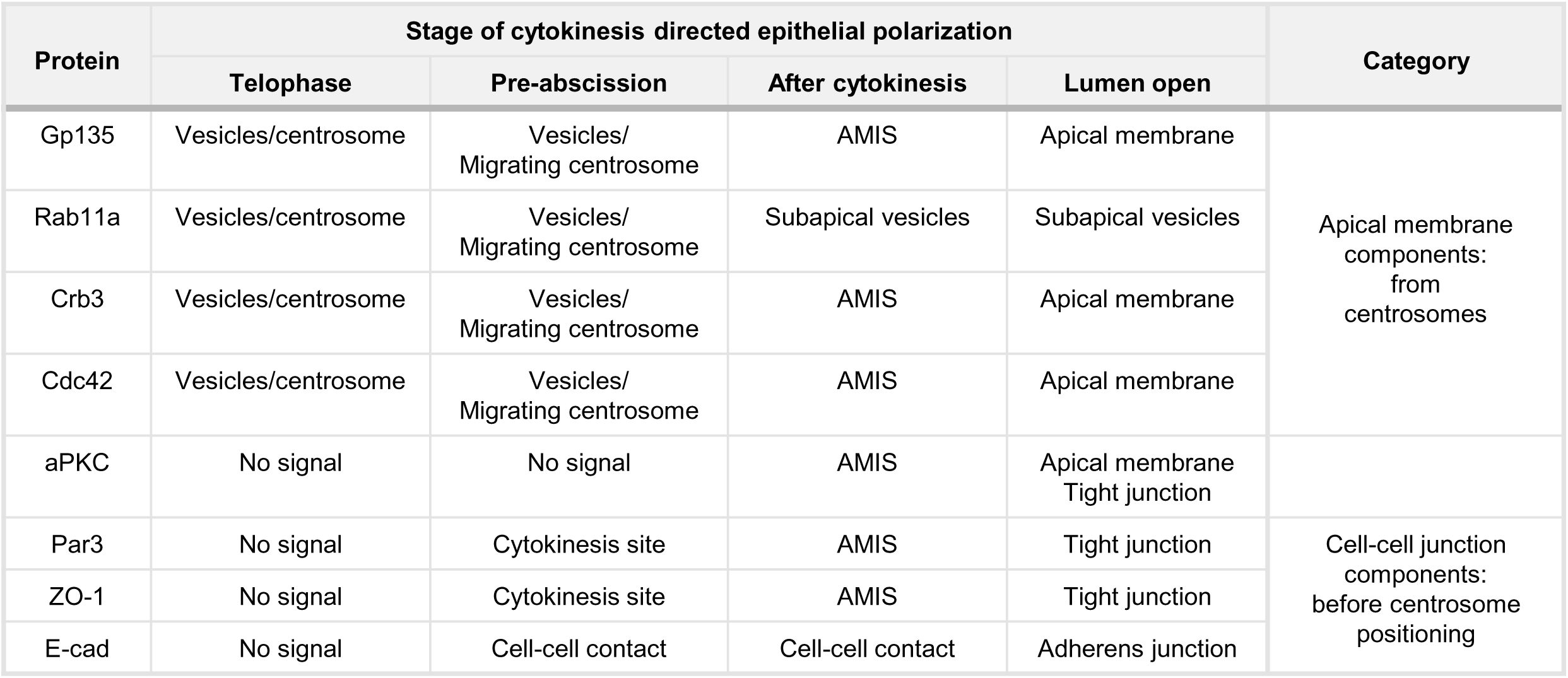
Distribution of trafficking and polarity proteins during cytokinesis-directed *de novo* epithelial polarization.

To establish causality between polarity marker Par3 and polarized transport, we generated Rab11a-KO cells (Figure 5—figure supplement 2C–E). In Matrigel culture, we found that the Par3 signal still appears first at the cytokinesis site and that the centrosome begins to migrate toward the center of the cell doublet in Rab11a-KO cells (Figure 5E). Similarly, the complete depletion of centrosomes in centrinone-treated p53-KO cells did not affect Par3 localization (Figure 5F). These data provide evidence in the reverse direction, suggesting that centrosome positioning and polarized vesicle trafficking are downstream of Par3 recruitment.

### Cytokinesis promotes the central migration of centrosomes during *de novo* epithelial polarization

Researchers have demonstrated an association between cytokinesis and the *de novo* polarization of epithelial cells (Klinkert et al., 2016; Li et al., 2014; Mangan et al., 2016; Rathbun et al., 2020; Schluter et al., 2009; Wang et al., 2014). Based on our results, Par3, which first localizes to the cytokinetic bridge, affects the centrosome positioning in Matrigel culture of MDCK cells. Here, we prevented cell division or changed culture conditions, which may affect the Par3 distribution in cells, to shed light on the relationship among the centrosome position, polarity regulator, and apical membrane components.

We observed cell doublets in Matrigel formed through cell-cell aggregation, not cell division (Figure 6—figure supplement 1A, see Materials and methods). Live-cell imaging showed that Gp135 is enriched on the membrane facing outward and that centrosomes remain stationary for several hours (Figure 6A). In early aggregated cell doublets, Gp135 polarity is inverted, consistent with previous studies (Bisi et al., 2020; Bryant et al., 2010; Bryant et al., 2014; Chou et al., 2016; Martin-Belmonte et al., 2007; Roman-Fernandez et al., 2018). We further performed immunostaining to examine polarity molecules. We found that Par3 was recruited to the cell-cell interface in the aggregated doublets (Figure 6B). Rab11a-positive recycling endosomes were concentrated around the centrosome (Figure 6C). Quantitative measurement *θ_N-C_* revealed that, although centrosomes are located between the two nuclei in the doublet generated by cytokinesis (Ctrl, Figure 6D, E), centrosomes are widely distributed across both nuclei in the aggregated doublets (+Aphi, Figure 6D, E). The results of the aggregated doublets indicate that the relationships among Gp135-associated apical membrane formation, Par3 location, and centrosome positioning are distinct from that of cytokinesis-induced *de novo* epithelial polarization.

**Figure 6:**
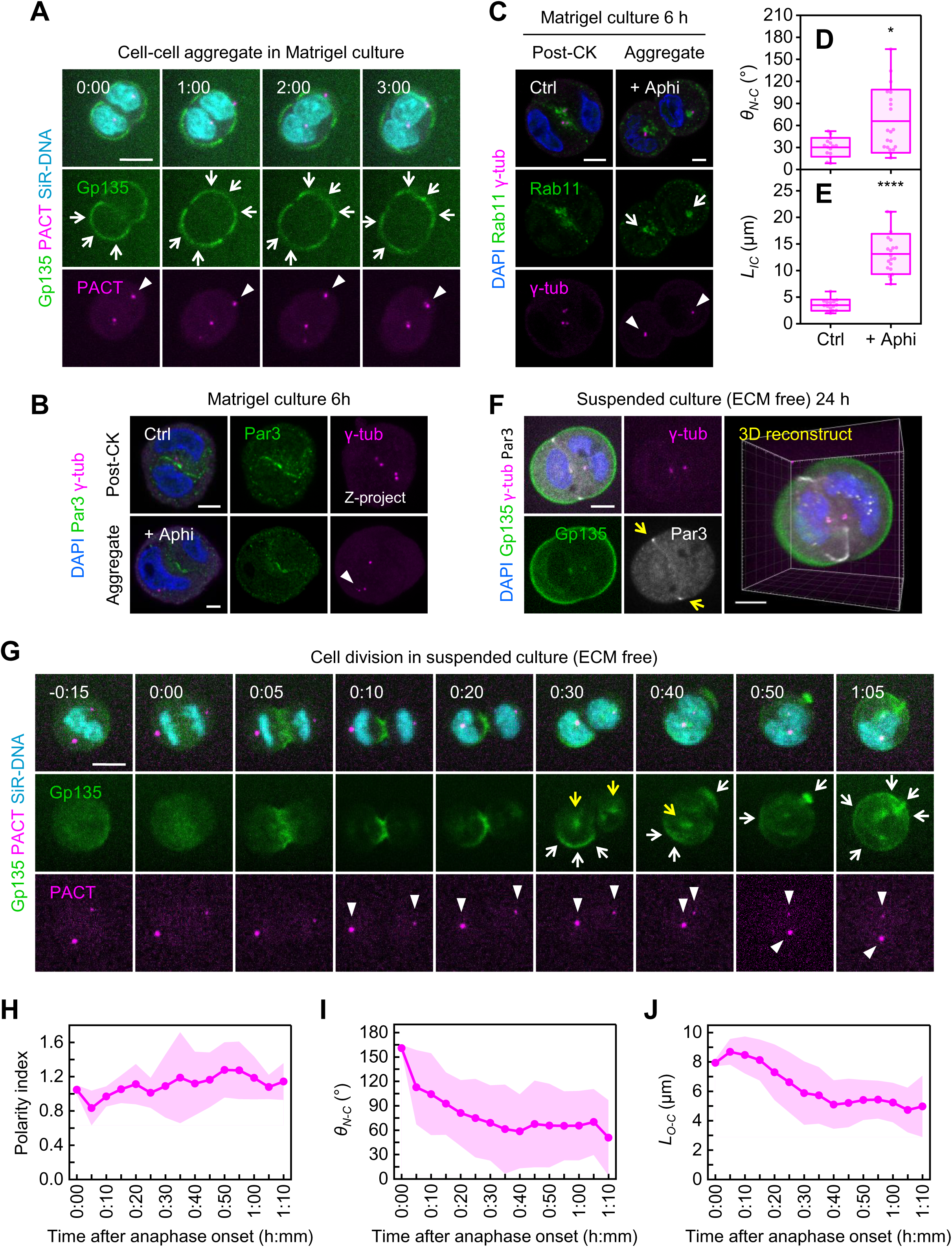
Centrosome migration in suspended 3D culture during cytokinetic pre-abscission. (A) Time-lapse snapshots of MDCK cells expressing EGFP-Gp135 (green) and PACT-mKO1 (magenta, centrosome marker) in Matrigel. Nuclei were labeled with SiR-DNA (cyan) before live imaging. Z-projection images of an aggregated cell doublet are shown. Arrows indicate Gp135 on the outer surface of the cell aggregate. Arrowheads point to mislocalized centrosomes that did not move over 3 h. Time stamps show hours and minutes, with 0:00 set at the beginning of live cell imaging. Scale bar: 10 μm. (B, E) MDCK cells, with or without aphidicolin (Aphi) synchronization, were cultured in Matrigel for 6 h with labeled markers: Rab11a (B, green), Par3 (E, green), γ-tubulin (magenta), and DAPI for nuclei (blue). Single confocal sections are shown. Two types of cell doublets, formed by cell division (post cytokinesis [CK]) or aggregation, were compared. Arrows indicate Gp135 clusters around centrosomes. Arrowheads point to mislocalized centrosomes. Scale bar: 5 μm. (C–D) Boxplots of centrosome positions, represented by *θ_N-C_* (C) and *L_IC_* (D), in cell doublets (n = 15 [unsynchronized control], 20 [Aphi-synchronized cells]). Cells were analyzed in three independent experiments. Statistical analyses used an unpaired two-tailed Mann–Whitney U test (*p<0.05, ****p<0.0001). Midlines and boxes show the mean ± SD, with whiskers indicating minimum and maximum values. (F) MDCK cells were cultured in low-attachment microwells for 24 h with labeled markers: Gp135 (green), γ-tubulin (magenta), Par3 (white), and DAPI for nuclei (blue). A single confocal section through the middle of a cell doublet is shown. Yellow arrows indicate the Par3 signal at the edge of cell-cell contacts. The 3D reconstruction reveals Par3 forming a ring around the edge between cell doublets. Scale bar: 10 μm. (G) Time-lapse snapshots of MDCK cells expressing EGFP-Gp135 (green) and PACT-mKO1 (magenta, centrosome marker) in low-attachment microwells. Nuclei were labeled with SiR-DNA (cyan) before live imaging. Z-projection images of a dividing cell are shown. White and yellow arrows indicate Gp135 on the outer surface of the cell doublet and around the centrosome, respectively. Arrowheads point to the centrosome position. Time stamps show hours and minutes, with 0:00 set at the first frame of anaphase onset. Scale bar: 10 μm. (H–J) Change in polarity index (H), *θ_N-C_* (I), and *L_O-C_* (J) over time. Each data point represents the average at a given time (three cells in low-attachment microwells from three independent experiments). The lines show the means, and the shaded regions indicate SD values.

We next create inverted cysts by culturing suspended single cells in low-adhesion microwells (Figure 6—figure supplement 1B) (Huang et al., 2020; Yu et al., 2005); in these conditions, the normal cell cycle progresses, but cells exhibit an inverted Gp135 polarity from cells in Matrigel culture. Immunostaining shows that Par3 proteins form a ring at the edge of the cell-cell interface (Figure 6F), distinct from the Par3 patch observed at the center of the cell-cell interface for cells in Matrigel (Figure 5—figure supplement 1H).

To observe the establishment of inverted polarity, cells were synchronized at prometaphase using nocodazole, then released and cultured in low-adhesion microwells, followed by live-cell imaging (Figure 6—figure supplement 1C). Before and at the onset of anaphase (Figure 6G, -0:15–0:00), the centrosomes form two spindle poles at two sides of the nuclei. At this time point, there is a dispersed Gp135 signal inside the cytoplasm. When cleavage furrow ingression occurs (Figure 6G, 0:05–0:20), Gp135 is concentrated at the furrow membrane between the two daughter cells. As cytokinesis progresses, Par3 first appears at the cell-cell interface at the mitotic cleavage site during the cytokinetic pre-abscission stage, similar to what is observed in Matrigel culture (Figure 6—figure supplement 1D). Gp135 dissociates from the middle membrane and becomes concentrated around the migrating centrosomes and external membranes (Figure 6G, 0:30, white and yellow arrows, Figure 6—figure supplement 1E). While Gp135 spreads toward the outside membrane, the centrosomes migrate toward the center of the cell doublets (Figure 6G, 0:40–1:05, and H-J). Finally, Gp135 is concentrated at the membranes, corresponding to locations where cells are not in contact, and forms a polarization opposite to that of cysts in Matrigel. Taken together, our results identify an interesting condition in which the polarity indicated by the centrosome and nucleus is opposite to the apical-basal polarity. In this case, the apical side is marked by Gp135, whereas the opposite pole is basal and in contact with adjacent cells. Furthermore, cytokinesis drives centrosome migration toward the center of the cell doublets, even in the absence of ECM and under conditions of inverted polarity.

Finally, we investigated the movement of centrosomes at the cytokinetic pre-abscission stage in highly polarized cysts (Figure 6—figure supplement 2A) and in an epithelial sheet (Figure 6—figure supplement 2B, C), of which both have apical-localized cytokinetic bridges. Both live cell imaging and immunostaining reveal that, in telophase cells, centrosomes are initially localized laterally at the mid-plane of the apical–basal axis and subsequently migrate apically to the site of the cytokinetic bridge during the pre-abscission stage (Figure 6—figure supplement 2A–C). Consistent with *de novo* epithelial polarization, we found that, when the apical membrane has not yet formed, the centrosomes move toward the center of cell doublets where the cytokinetic bridge is located (Figure 6—figure supplement 2A–D). Taken together, these findings indicate that the centrosome position is strongly associated with the cytokinesis site but not with the apical membrane.

## Discussion

Because the vector from the nucleus to the centrosome is often observed aligning with the axis of other polarization markers in various cells, the centrosome has long been hypothesized to play a crucial role in regulating cell polarization (Bornens, 2012; Burakov & Nadezhdina, 2020; Tang & Marshall, 2012). Based on the significant movement of centrosomes during polarization derived from cell division, as well as the dependence of polarity molecules and apical membrane location on culture conditions, we designed a series of experiments to test the role of the centrosome. Our findings, which highlight the correlation between centrosome positioning and cell polarization, as well as the impact of altered centrosome function on the distribution of polarity markers, are summarized in Figure 7.

**Figure 7:**
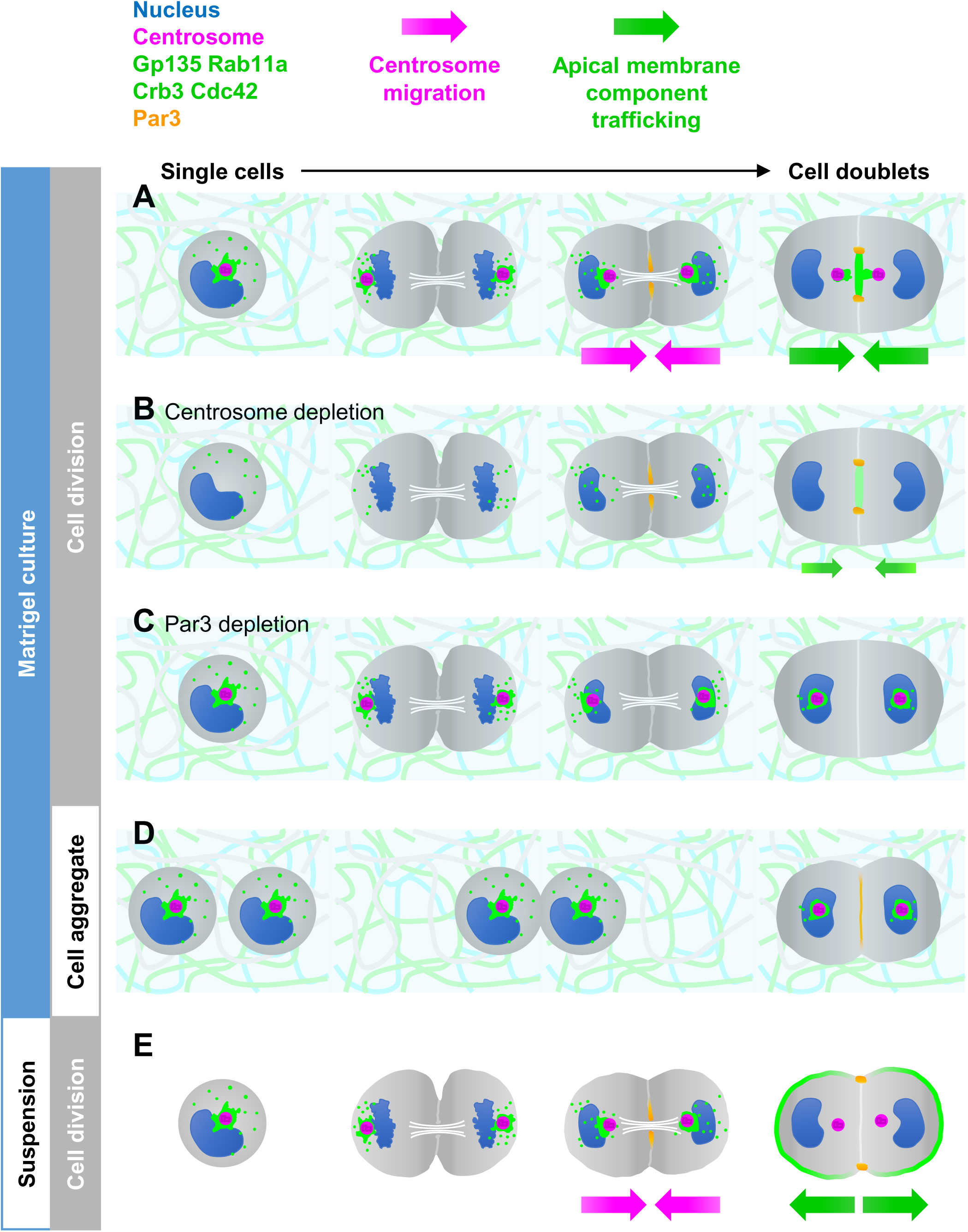
Summary of centrosome migration, apical membrane component trafficking, and Par3 recruitment in different experiments and culture conditions. (A) In Matrigel culture, centrosomes move directionally toward AMIS following cell division (magenta arrow). Apical membrane components like Gp135, Crb3, and Cdc42 follow the centrosomes (green arrow). Par3 first emerges at the cytokinesis site, which regulates centrosome positioning and polarized vesicle trafficking during polarization. (B) Loss of centrosomes diminishes the effectiveness of apical membrane component trafficking (smaller green arrow), yet Par3 still localizes to the cytokinetic bridge. Additionally, centrosome loss mainly impacts the initial stages of epithelial polarization rather than the later stages of lumen formation. (C) Loss of Par3 results in randomized centrosome positioning, obstructs polarized trafficking, and traps apical membrane components around mislocalized centrosomes. (D) Two-cell aggregates without cell division cannot effectively guide centrosome migration and polarized vesicle trafficking. Despite Par3 being recruited to the cell-cell interface, apical membrane components remain trapped around mislocalized centrosomes. (E) MDCK cells suspended in the ECM-free condition. Centrosomes migrate to the center of cell doublets during cytokinesis (magenta arrow), while Apical membrane components transport in the opposite direction (green arrow). After mitosis, Par3 exhibits a pattern distinct from that observed in Matrigel culture.

In Matrigel culture, where *de novo* epithelial polarity develops, our findings show that apical membrane components, including Gp135, Crb3, and Cdc42, accumulate around the centrosomes and follow the rapidly moving centrosomes to the AMIS during cytokinesis (Figure 1 and 2). Par3 first appears at the cytokinetic bridge, earlier than the positioning of the centrosome and the targeting of apical membrane components (Figure 5 — figure supplement 1H). The aligned direction of centrosome positioning, apical membrane protein trafficking, and Par3 localization significantly enhance the efficiency of polarization (Figure 7A). The centrosome depletion experiments indicate that loss of centrosomes still allows the formation of AMIS and lumen structures (Figure 3 and 5F), similar to their dispensable roles in cell division or spindle microtubule formation (Meitinger et al., 2016; Watanabe et al., 2020; Wong et al., 2015). Under p53-KO conditions, centrosome-depleted MDCK cells can form cysts with a single central lumen at a slower rate. However, due to decreased trafficking efficiency, the protein levels on the apical membrane are reduced (Figure 7B). Previous studies have debated the role of centrosomes in various types of cell polarization (de Anda et al., 2010; Martin et al., 2018; Schmoranzer et al., 2009; Stiess et al., 2010; Wakida et al., 2010). We determine that while the centrosome is not essential, it has a synergistic effect on apical membrane establishment.

In addition to complete removal of the centrosome, we also disrupted specific substructures within the centrosome to investigate which components are responsible for regulating centrosome positioning, vesicle trafficking, and cell polarization. In particular, appendage structures and PCM proteins that link centrosomes to microtubules are considered prime candidates, as microtubules are thought to generate pushing and pulling forces that facilitate centrosome positioning (Hooikaas et al., 2020; Kapitein et al., 2005; Yi et al., 2013). However, we did not identify a single centrosomal molecule whose disruption alone significantly affected centrosome positioning (Figure 4). These findings led us to further investigate whether centrosome-nucleating microtubules are essential for proper centrosome positioning and polarized vesicle trafficking. By competitively displacing γ-tubulin from centrosomes, we were able to produce centrioles devoid of associated γ-tubulin. Due to the displacement of γ-tubulin, these centrioles were unable to nucleate microtubules, demonstrating the effective elimination of centrosomal microtubules. Interestingly, despite the loss of centrosomal microtubules, centrioles were still localized to the apical region of Matrigel-cultured MDCK cells (Figure 4—figure supplement 3). These findings challenge the previously proposed model that centrosomal microtubules are required to actively pull centrosomes to a specific cellular position. Instead, they suggest that additional mechanisms contribute to centrosome positioning. For example, recent studies propose that motor proteins can transport centrosomes as cargo along non-centrosomal microtubules (Hannaford et al., 2022; Hannaford & Rusan, 2024), while others implicate that the actin cytoskeleton is involved in regulating centrosome positioning (Jimenez et al., 2021; Yamamoto et al., 2022).

Among all our experiments involving molecular perturbations to elucidate relationships between different polarity features, we have discovered that Par3 plays a pivotal role in upstream signaling during cytokinesis-induced polarization, which affects centrosome positioning and polarized trafficking (Figure 7C). Par3 has been reported to regulate centrosome position in micropattern-cultured MCF10A cells and in the intestine of *C. elegans* (Burute et al., 2017; Feldman & Priess, 2012) and has also been shown to affect centrosome position through interaction with dynein in migrating NIH 3T3 cells (Schmoranzer et al., 2009). However, Par3 is not directly localized to the centrosome, but rather to the cell cortex or cell-cell junctions. Therefore, Par3 is likely to regulate centrosome positioning through some intermediate molecules or mechanisms. One possibility is that the interaction between Par3 and dynein anchors dynein to the cell cortex, thereby generating a pulling force along the microtubules to position the centrosomes, as reported in NIH 3T3 cells and *C. elegans* (Feldman & Priess, 2012; Schmoranzer et al., 2009). However, our immunofluorescence analysis did not reveal colocalization of dynein with Par3 at the cytokinetic bridge. The specific mechanism is still unclear and requires further investigation. Pervious study have also suggested that Par3 functions as an exocyst receptor in epithelial cells, mediating the targeting of delivered proteins to specific domains of the plasma membrane through interaction with Sec8 and Exo70, two exocyst components (Ahmed & Macara, 2017). Depletion of Par3 leads to membrane proteins and Rab11a residing in the cytoplasm (Ahmed & Macara, 2017; Horikoshi et al., 2009). Consistent with these findings, we found that Rab11a-positive vesicles sometimes pause during delivery and become trapped around mislocalized centrosomes in Par3-depleted MDCK cells (Figure 5A and Figure 5—figure supplement 1G). Taken together, our findings support the concept that Par3 participates in centrosome positioning and mediates the transportation of proteins on the Rab11a-positive endosomes to AMIS during cytokinesis-induced polarization. However, cultured conditions influence cellular polarization preferences. For instance, when cells aggregate in Matrigel, we observe recruitment of Par3 to the cell-cell interface, despite the absence of a division site to specify its location, but both centrosome positioning and Gp135 transport are disrupted (Figure 6A-E and 7D). It is necessary to determine the differences in the function of Par3 between cell aggregates and during cytokinesis, such as whether it also interacts with dynein and exocysts in these cell aggregates.

When MDCK cells are suspended in ECM-free conditions, they form inverted cysts after division, with Gp135 facing outward, while Par3 forms a ring at the edge of the cell-cell contact surface (Figure 6F-J). Intriguingly, the vector from the nucleus to the centrosome is opposite from the direction of the apical membrane (Figure 7E). Notably, we found that centrosomes do not constantly localize near the apical surface. These different cultured conditions help us dissect the process of epithelial polarization and the relationships between different polarity indicators. During epithelial polarization, centrosome positioning, apical membrane component trafficking, and the recruitment of the polarity regulator Par3 appear to be executed through distinct mechanisms. It is known that apical membrane components undergo endocytosis and are transported by the Rab11a cascade to target in the opposite direction of the ECM contact side (Jewett & Prekeris, 2018; Yu et al., 2005). However, the mechanisms for centrosome positioning and Par3 recruitment remain unclear. Recent studies have shown that the cortical actin flows in *C. elegans* blastomeres attracts the enrichment of Par3 (Ng et al., 2023), and that the acto-myosin network plays a role in the centrosome off-centering process in enucleated cells (Jimenez et al., 2021; Yamamoto et al., 2022). Therefore, the dynamics of the actomyosin network may be crucial for both Par3 recruitment and centrosome migration, as it undergoes significant rearrangements during cytokinesis. During the pre-abscission stage, the actomyosin network loses its symmetric distribution and a contractile ring forms at the center of the cell doublet, while the amount of actin decreases at the cell periphery (Figure 5—figure supplement 2A). These events during cytokinesis may promote Par3 recruitment and trigger centrosome migration. Subsequently, a dense actin network remains at the center of the cell doublet, contributing to the formation of the apical actin network during the post-cytokinesis stage. In Matrigel cultures, ECM signals and the actin network are aligned in the same direction, synergistically enhancing the efficiency of epithelial polarization. In cell-aggregated conditions, cells have not yet divided and the actin network does not show dynamic changes. This lack of remodeling may prevent centrosome movement, resulting in a reduced efficiency of epithelial cell polarization.

Although our investigation primarily focused on the 3D culture of epithelial cells *in vitro*, the insights gleaned from this study may have implications for understanding polarity establishment in various epithelial-like tissues *in vivo*.

## Materials and methods

### 3D cell culture in Matrigel and drug treatment

MDCK (NBL-2) cells (Bioresource Collection and Research Center, Taiwan) were tested for mycoplasma, cultured in minimum essential medium (MEM) with glucose, 2 mM L-glutamine, and 10% fetal bovine serum, and supplemented with 0.1 mM non-essential amino acids and 1 mM sodium pyruvate, penicillin, and streptomycin. We conducted 3D cell culture assays for cyst formation using a previously described method (Lee et al., 2007; Martin-Belmonte et al., 2007). We trypsinized 70% of confluent cells to a single cell suspension of 4 × 10^4^ cells/mL in complete MEM containing 2% growth factor reduced Matrigel (356231, Corning, Glendale, AZ). We plated cells in 2% Matrigel medium (300 μL) in 8-well cover-glass chamber slides (80827, ibidi, Gräfelfing, Germany) pre-coated with 8 μL of 100% Matrigel. The cells were incubated for the indicated period, and the medium was changed every 2 days.

For the centrosome depletion assay, we pre-treated MDCK WT and p53-KO cells with 300 nM centrinone (LCR-263, HY-18682, MedChemExpress, Monmouth Junction, NJ) (Wong et al., 2015) for 3 days on a petri dish to achieve >80% of cells without centrosomes. The centrosome-depleted cells were then trypsinized to a single-cell suspension and plated into a 3D culture in the continuous presence of 300 nM centrinone.

To assess cell aggregates, we treated MDCK cells with aphidicolin (Aphi, 2.5 μg/ml) for 15 hours, arresting almost all cells in the S phase. Subsequently, we released these synchronized cells from the S phase and simultaneously cultured them in Matrigel. Based on past experience, MDCK cells require approximately 8 hours to progress from the S phase to the M phase. Therefore, we observed cell doublets at 6 hours after release to ensure that these cell doublets were formed through aggregation rather than cell division.

To enrich mitotic cells in suspension culture, MDCK cells were treated with nocodazole (100 ng/mL) for 4 hours to arrest cells predominantly at the prophase stage. Cells were then trypsinized and released from the nocodazole block, and immediately transferred to low-attachment culture dishes to maintain suspension conditions. Based on previous observations, cell doublets typically appeared ∼1 hour after release, corresponding to the telophase to pre-abscission stages.

### Antibodies

We used rabbit polyclonal antibodies against Gp135 (IF: 1:400, a kind gift from Dr. Jou Tzuu-Shuh, NTU, Taiwan) (Lim et al., 2017), Rab11a (IF: 1:400, WB: 1:2000, 20229-1-AP, Proteintech, Rosemont, IL), aPKC (IF: 1:200, 610175, BD Biosciences, Franklin Lakes, NJ), CEP120 (IF: 1:1000, WB: 1:2000, human residues 639–986, described in our previous paper) (Lin et al., 2013), p53 (IF: 1:100, 9282, Cell Signaling Technology, Danvers, MA), p21 (WB: 1:2000, 10355-1-AP, Proteintech), CEP164 (IF: 1:400, NBP1-81445, NOVUS, Centennial, CO), PCNT (IF: 1:200, ab4448, Abcam, Cambridge, UK), Par3 (IF: 1:200, WB: 1:1000, 07-330, EMD Millipore, Burlington, MA), and beta-actin (IF: 1:200, A2066, Sigma-Aldrich, St. Louis, MO); rabbit monoclonal antibodies against E-cadherin (IF: 1:200, 24E10, #3195, Cell Signaling Technology); and mouse monoclonal antibodies against γ-tubulin (IF: 1:400, GTU-88, T6557, Sigma-Aldrich), acetylated tubulin (IF: 1:400, T6793, Sigma-Aldrich), Gp135 (IF: 1:10, the culture medium of hybridoma clone 3F2D8, a kind gift from Dr. Jou Tzuu-Shuh, NTU, Taiwan) (Lim et al., 2017), polyglutamylated tubulin (IF: 1:100, GT335, AG-20B-0020, Adipogen, San Diego, CA), ODF2 (IF: 1:400, H00004957-M01, NOVUS), α-tubulin (WB: 1:5000, DM1α, T9026, Sigma-Aldrich), CDC42 (WB: 1:1000, 610928, BD Biosciences), and ZO-1 (IF: 1:400, 33-9100, ThermoFisher Scientific, Waltham, MA).

For western blots, we used the following secondary antibodies: HRP-labeled goat anti-rabbit and mouse IgG (1:10000, NEF812 and NEF822, PerkinElmer, Waltham, MA). For immunofluorescence, we used Alexa Fluor 488, 561, and 647 conjugated goat antibodies against rabbit, mouse IgG1, and mouse IgG2b (1:400, all purchased from ThermoFisher Scientific) as secondary antibodies, together with Phalloidin-iFluor 647 reagent (1:1000, ab176759, Abcam) and 4’,6-diamidino-2-phenylindole (DAPI) (1:10000, D8417, Sigma-Aldrich).

### 3D immunofluorescence staining, image acquisition, and analysis

We stained 3D cell cultures according to a previously published protocol, with some modifications (Lee et al., 2007). We pre-fixed whole cultures in each well with 4% paraformaldehyde at room temperature for 10 min. If the staining included centrosome markers, we post-fixed the samples with pure cold methanol at -20°C for another 10 min. After fixation, cells were permeabilized with 0.25% Triton X-100 in phosphate-buffered saline (PBS). We performed primary antibody staining in 100 μL PBS plus 10% normal goat serum at 4°C for 48 h. After four 15 min washes in 250 μL PBS with 0.1% Tween 20 at room temperature, we added secondary antibodies with DAPI in 100 μL PBS plus 10% normal goat serum for another 48 hours at 4°C. Finally, we performed another four 15 min washes, followed by adding 100 μL mounting medium (VectaShield, Vector Laboratories, Newark, CA) to each well. We analyzed the samples on the LSM 880 confocal (Zeiss, Jena, Germany) using a Plan-Apochromat 63x oil immersion objective with a numerical aperture (NA) 1.4.

For fluorescence intensity measurements, confocal images were acquired with constant parameters (pinhole size, laser power, master gain, and z-stack interval) and introduced to Fiji. We applied the maximum intensity projection to project the z-stack. The oval selection tool was fixed on size to select the region of interest (ROI) (e.g., pericentrosomal region, cytokinetic bridge, cytosol, AMIS, or apical lumen) and to measure the mean fluorescence intensity of the ROI.

### Quantification of centrosome position

We considered the polarity index, *θ_N-C_*, *L_O-C_*, and *L_IC_* of cell doublets cultured in 3D conditions to assess centrosome positions. We obtained the centroid of each cell nucleus and centrosome, as well as the volume of the cell nucleus, using Imaris software (version 9.8.0, Bitplane, Belfast, UK). The cell nucleus radius, r_N_, was determined by dividing its volume by 4/3π and then taking the cube root. We calculated the polarity index from the nucleus–centrosome distance, *L_N-C_*, divided by the *r_N_*.

The nucleus–centrosome orientation (*θ_N-C_*) is represented by the angle between the line from the nucleus to the centrosome and the line from the nucleus to the other nucleus. The central points of each nucleus and centrosome were marked as three points (nuclei: n1 and n2; centrosome: c) in 3D space to give three x, y, z coordinates. We computed the angle between the vector of the nucleus to the centrosome and the vector of the nucleus to the other nucleus using the vector dot product formula.

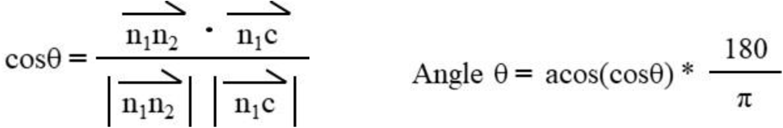

Centrosomes oriented within 90° from the nucleus–nucleus axis were considered to be correctly oriented toward the apical region.

The distance between the centrosome and apical membrane was represented by the distance from the origin point (the center point of the two nuclei) to the centrosome, *L_O-C_*, because the AMIS and apical lumen will form at the center of the cell doublet in Matrigel culture. We also calculated the distance using x, y, z coordinates in 3D space. A smaller distance indicates that the centrosome and apical membrane are closer. The distance between the centrosomes in each cell of a cell doublet is referred to as the inter-centrosomal distance, *L_IC_*.

### Lentiviral transductions and stable cell lines

We generated MDCK cells expressing EGFP-rabbit podocalyxin (Gp135, complementary DNA [cDNA] was a kind gift from K. Simons), EGFP-Crb3 (cDNA was obtained by reverse transcription polymerase chain reaction [RT-PCR] following previously described procedures) (Schluter et al., 2009), EGFP-Cdc42, PACT-mKO1, mCherry–C-CTD, and mCherry–N-gTBD via lentiviral transduction using the following protocol: MDCK cells were transduced with lentiviruses that package with the pLAS2w.Ppuro vector (RNAi Core, Taiwan) containing the relevant cDNA. The transduction processes were performed in complete MEM with 8 μg/mL polybrene. The transduced cells were sterile-sorted by a cell sorter (FACSJazz, BD Biosciences) for EGFP, mKO1, or mCherry. Fluorescence-positive cells were selected and expanded.

### 3D live cell imaging in Matrigel and analysis

We resuspended stable MDCK lines in a culture medium containing 2% Matrigel and plated the cells in a rose chamber with the cover-glass bottom pre-coated by 100% Matrigel. We allowed the cells to remain in the incubator for 2 hours to ensure that the cells fell and were fully adherent on the Matrigel. We labeled the nucleus in live cells with a far-red fluorescence probe, SiR-DNA (500 nM, SC007 Spirochrome, Cytoskeleton Inc., Denver, CO), in phenol red-free medium containing 2% Matrigel.

After a 1 hour incubation, we performed live cell imaging on a 40x NA1.1 water immersion objective (Leica, Wetzlar, Germany) on an inverted microscope (DMI 6000, Leica) with a confocal scan head (CSU22 Spinning Disk, Yokogawa Electric, Tokyo, Japan), a laser merge module containing 405-, 491-, 561-, and 642-nm laser lines (ILE-400, Andor, Belfast, UK), a DV885 EMCCD (Andor), an MS-2000-XYZ motorized stage with a piezo top plate (ASI, Eugene, OR), and a stage-top incubator set to 37°C under 5% CO_2_. The microscope apparatus was controlled by MetaMorph 7.7.2 (Molecular Devices, San Jose, CA). We set the time interval to 5 min and z steps to 0.5 μm.

We collected images of mitotic cells to track centrosome migration during *de novo* epithelial polarization. These images were imported to Imaris software (version 9.8.0, Bitplane) for 3D reconstruction and analysis. We marked the centroid of each nucleus and centrosome as x, y, z coordinates in 3D space for each time point. We calculated the polarity index, *θ_N-C_*, and *L_O-C_* from the first frame of anaphase onset.

To measure the fluorescence intensity of EGFP-Gp135 on the middle membrane of the 3D cell doublet, we measured the intensity profile of EGFP-Gp135 along the line connecting the two nucleus spots. Then, we recorded the average value of the five center pixels representing the region of the middle membrane from the first frame of anaphase onset.

### CRISPR/Cas9 knockouts

To generate MDCK lines lacking either p53, CEP164, ODF2, CEP120, PCNT, or Rab11a, we used CRISPR/Cas9 technology. Because CEP120-null cells cannot survive in the presence of WT p53 (Tsai et al., 2019), we performed a knock-out of CEP120 in MDCK p53-null cells. We transfected MDCK cells using a Lipofectamine 2000 transfection reagent (11668019, ThermoFisher Scientific) with the pSpCas9(BB)-2A-GFP (PX458) vector (Ran et al., 2013) (648138, Addgene, Watertown, MA) bearing the appropriate targeting sequence: *p53* #1 (+): 5’-TCCCAGAGAGCGTCGTGAAC-3’; *p53* #2 (-): 5’-ACTTGGCTGTCCCGGAATGC-3’; *CEP164* #1 (-): 5’-GTCCCACATGGACTGCCCGT-3’; *CEP164* #2 (+): 5’-AACTTGGTGATTCAAGAGCG-3’; *ODF2* #1 (+): 5’-CCTCATGTCCAAGCTGGTAG-3’; *ODF2* #2 (+): 5’-ATCGGGAAGCTGAAGACGGT-3’; *CEP120* #1 (+): 5’-ACAGTTGGCTACCGACCCTG-3’; *PCNT* #1 (+): 5’-TCGCCAGGGAGCAGCATGCG-3’; *Rab11a* #1 (-): 5’-GCGACGACGAGTACGACTAT-3’; *Rab11a* #2 (-): 5’-AGCTATAACATCAGCGTAAG-3’

Cells were sterile-sorted by a cell sorter (FACSJazz, BD Biosciences) for GFP. Fluorescence-positive cells were serially diluted to pick single colonies and were expanded to pure clones. Genomic DNAs isolated from each clone were subjected to PCR and sequencing for verification. We examined protein expression levels in each clone via immunofluorescence microscopy or western blotting.

### RNA interference

We achieved RNA interference (RNAi) using pLKO.1-EGFP (TRC011, RNAi Core, Taiwan) lentiviral vector. RNAi target sequences were based on previously published reports (Bryant et al., 2010; Horikoshi et al., 2009; Roland et al., 2011; Schluter et al., 2009) for canines, as follows: Rab11a #1: 5’-AAGGCACAGATATGGGACACG-3’; Rab11a #2: 5’-ATGGTTTGTCATTCATTGAGA-3’; Sec15a #1: 5’-AAGAGGATGAGAATGAAGAGG-3’; Sec15a #2: 5’-AAGCACGGGTCATGATAGTTT-3’; Par3 #1: 5’-GACAGACTGGTAGCAGTGT-3’; Par3 #2: 5’-AGGATAAAGCTGGCAAAGA-3’; MyoVb #1: 5’-AAGGTGGAGTATCTTTCAGAT-3’; MyoVb #2: 5’-AACGGGTAACTGTGGCCTTTA-3’; Cdc42 #1: 5’-GCGATGGTGCCGTTGGTAA-3’; Cdc42 #2: 5’-GATTACGACCGCTGAGTTA-3’; Crb3: 5’-GCTTAAGAGTAGAAGGGAA-3’

We verified knockdown by western blot or quantitative real-time PCR procedures normalized to GAPDH expression. The designed primer sequences were as follows: Sec15a Fwd: 5’-GTCAGCCTGCCAGCATCTGT-3’; Sec15a Rev: 5’-CTGCTGAACAGCTCCCATGC-3’; MyoVb Fwd: 5’-CAAGTTCCCACTGGTGGCTG-3’; MyoVb Rev: 5’-GAAGACGAGCCCTTCCCAGA-3’; Crb3 Fwd: 5’-GAGAACGGCACCATTACACC-3’; Crb3 Rev: 5’-GAGGGAGAAGACCACAATGAT-3’;GAPDH Fwd: 5’-AGTCAAGGCTGAGAACGGGAAACT-3’; GAPDH Rev: 5’-TGTTTGTGATGGGCGTGAACCATG-3’

### 3D live cell imaging in ECM-free conditions

Following a previously published protocol (Huang et al., 2020), we generated multiarray spherical microwells for an ECM-free suspension culture. In brief, 100 μL polyacrylamide (PA) solution containing 7% acrylamide monomer (Bio-Rad, Hercules, CA), 0.35% bisacrylamide crosslinker (Bio-Rad), 0.15% N, N, N’, N’-tetramethylethylenediamine catalyst (Sigma-Aldrich), and 0.5% ammonium persulfate (Sigma-Aldrich) was sandwiched between an epoxy mold and 3-(trimethoxysilyl) propyl methacrylate silanized cover glass (22 × 22 mm). Waiting for the complete gelation of PA, we carefully separated the cover-glass-bound microwell substrate from the epoxy template. To avoid cell adhesion, we conjugated poly-L-lysine/polyethylene glycol to the PA substrate using the bi-functional crosslinker sulfo-SANPAH (ThermoFisher Scientific). We sterilized low-attachment microwell substrates with antibiotics and then seeded a single-cell suspension of MDCK stable lines onto the substrate surface. After cells fell into the microwells, we rinsed the PA substrate with the culture medium to remove excess cells outside the microwells. The samples were assembled into a rose chamber and imaged by a CSU22 spinning disk confocal microscope, as described above, for 3D live cell imaging in Matrigel.

### Transwell-cultured 2D epithelial sheet

To grow polarized epithelial cell monolayers suitable for observation under an inverted microscope, we cultured MDCK cells on the outer surface of Transwell inserts (3470, Corning). Typically, 50 μL of a suspension containing 4.25 × 10^5^ cells/mL was seeded into each 6.5-mm Transwell insert. Cells were cultured for 3 days before immunofluorescence staining or live cell imaging. We mounted Transwell inserts in a custom-made glass-bottom chamber for confocal imaging. For live cell imaging, we stained epithelial cell monolayers with SiR-tubulin (500 nM, SC002, Spirochrome) and monitored the cells using LSM 980 confocal microscopy (Zeiss).

### Statistical analysis

Statistical analyses utilized GraphPad Prism 9 and OriginPro 2021, employing the Mann-Whitney U test and one-way and two-way analysis of variance (ANOVA) with multiple comparisons. Details on statistical tests and sample sizes are provided in the figure legends. n denotes the sample size, and the boxplot boxes indicate the standard deviation (SD).

## Conflict of interest

The authors declare no conflicts of interest.

## Author contributions

Conceptualization: P.-K. Wang and T. K. Tang; Investigation: P.-K. Wang; Methodology: P.-K. Wang and K.-H. Lin; Resources: P.-K. Wang and K.-H. Lin; Writing – Original Draft: P.-K. Wang, K.-H. Lin and T. K. Tang; Writing – Review & Editing: P.-K. Wang, K.-H. Lin and T. K. Tang.

## Ethical approval

N/A

## Funding

This work was supported by grants from Academia Sinica (AS-IA-109-L04) and the National Science and Technology Council, Taiwan (NSTC 112-2326-B001-010).

## Acknowledgments

We thank the DNA Sequencing Core Facility (IBMS, AS-CFII-113-A12), Flow Cytometry Core Facility (IBMS, AS-CFII-111-212), and Light Microscopy Core Facility of the Institute of Biomedical Sciences, Academia Sinica, for their valuable assistance. We also thank Dr. Tzuu-Shuh Jou (NTU, Taiwan) for the Gp135 antibody and Dr. Kai Simons (MPI-CBG, Germany) for the EGFP-rabbit podocalyxin cDNA.

## Legends to Supplemental Figures

**Figure 1—figure supplement 1:**
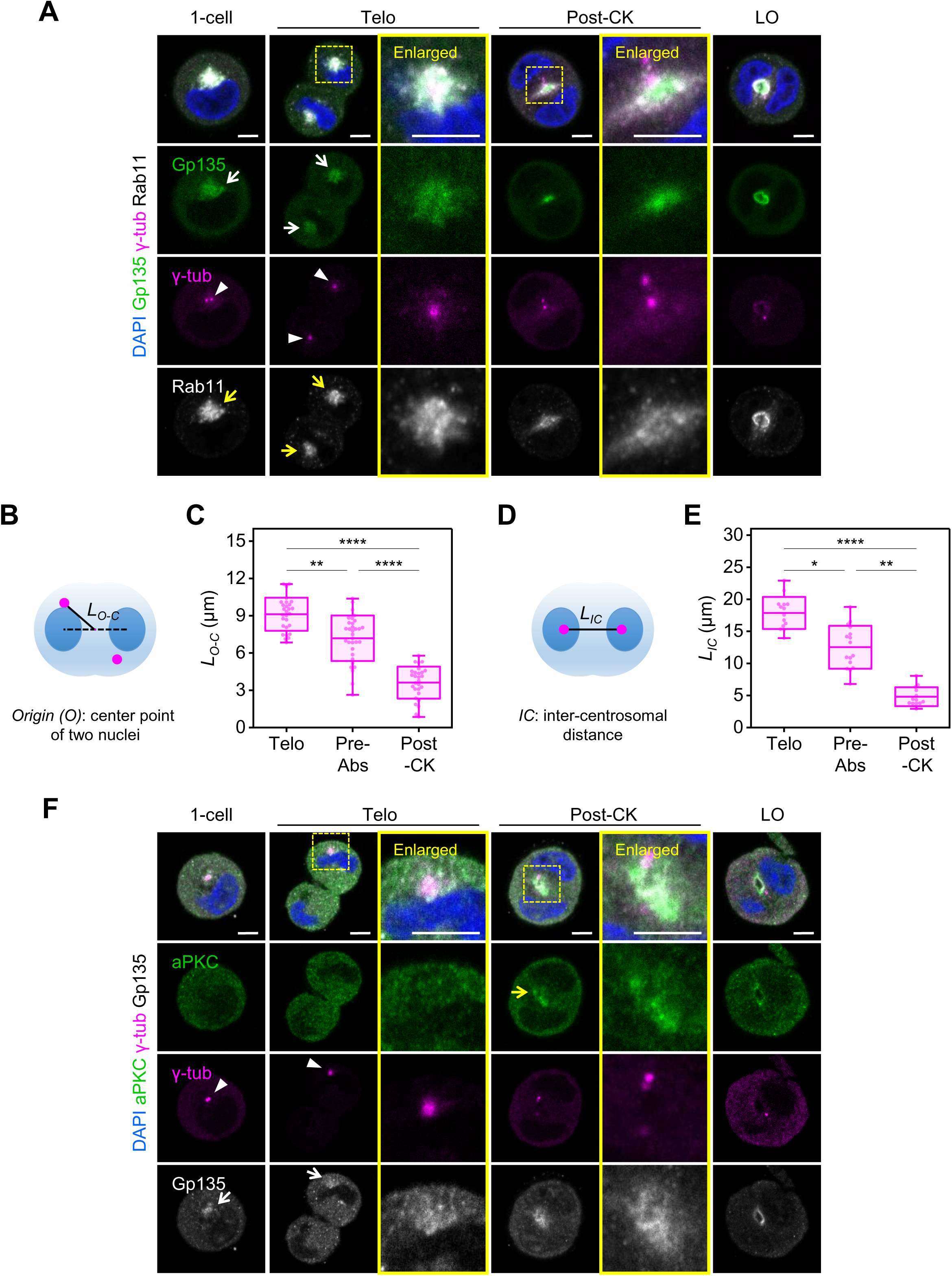
Distribution of Rab11a, Gp135, and aPKC proteins and centrosome position during *de novo* epithelial polarization. (A, F) Single MDCK cells were cultured in Matrigel for 12 h with labeled markers: Gp135 (green), γ-tubulin (magenta), and Rab11a (white) in (A), and aPKC (green), γ-tubulin (magenta), and Gp135 (white) in (F). DAPI staining was applied for nuclei (blue). Single confocal sections are shown through the middle of the cells. The centrosome region during telophase (Telo) and the AMIS region in post-cytokinesis (Post-CK) cells, as shown in the yellow box, are enlarged. Yellow arrows indicate Rab11a and aPKC around centrosomes or at the AMIS. White arrows indicate Gp135. Arrowheads point to the centrosome position. LO: lumen open. Scale bar: 5 μm. (B, D) Illustration showing *L_O-C_* (origin to centrosome distance) and *L_IC_* (inter-centrosomal distance) for cell doublets. The center point between the two nuclei is taken as the origin (O). (C, E) Boxplots depicting centrosome positions at different stages, shown by *L_O-C_* (C) and *L_IC_* (E). We analyzed >15 cells per measurement in three independent experiments. Statistical analyses used one-way ANOVA and Dunn’s multiple comparisons (*p<0.05, **p<0.01, ****p<0.0001). Midlines and boxes show the mean ± SD, with whiskers indicating minimum and maximum values.

**Figure 1—figure supplement 2:**
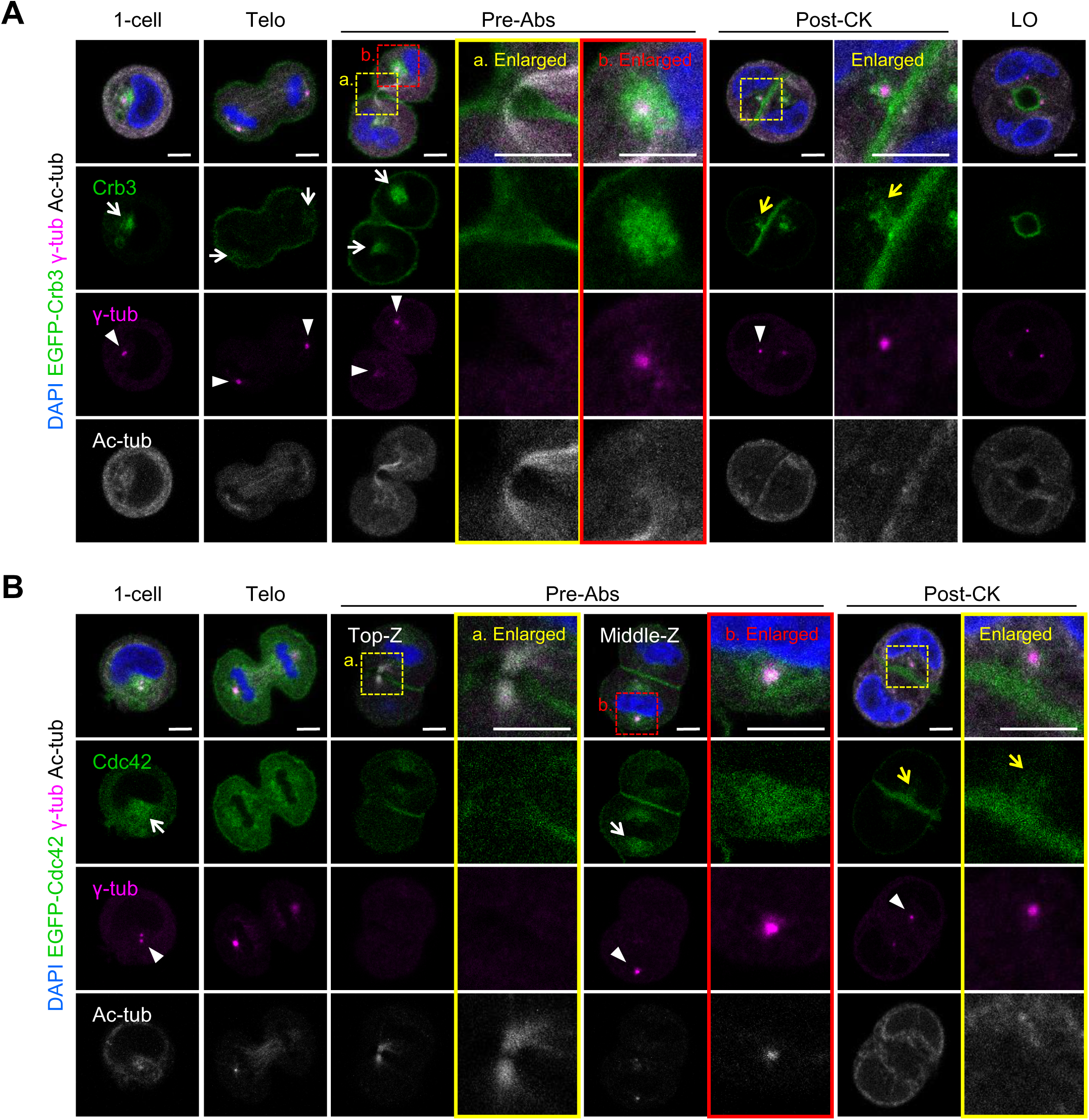
Clustering of Crb3 and Cdc42 around centrosomes during *de novo* epithelial polarization. (A–B) Single MDCK cells stably expressing EGFP-Crb3 (A, green) or EGFP-Cdc42 (B, green) were cultured in Matrigel for 12 h with labeled markers: γ-tubulin (magenta), acetyl-tubulin (white), and DAPI for nuclei (blue). Single confocal sections through the middle of cells are shown. The centrosome, cytokinesis (CK) bridge, and AMIS region are enlarged from the boxed area in pre-abscission (Pre-Abs) and post-CK cells. White arrows indicate EGFP-Crb3 (A) or EGFP-Cdc42 (B) around centrosomes. Arrowheads point to the centrosome position. Yellow arrows indicate the transport of Crb3 and Cdc42 from the centrosome to the mid-membrane. Scale bar: 5 μm.

**Figure 2—figure supplement 1:**
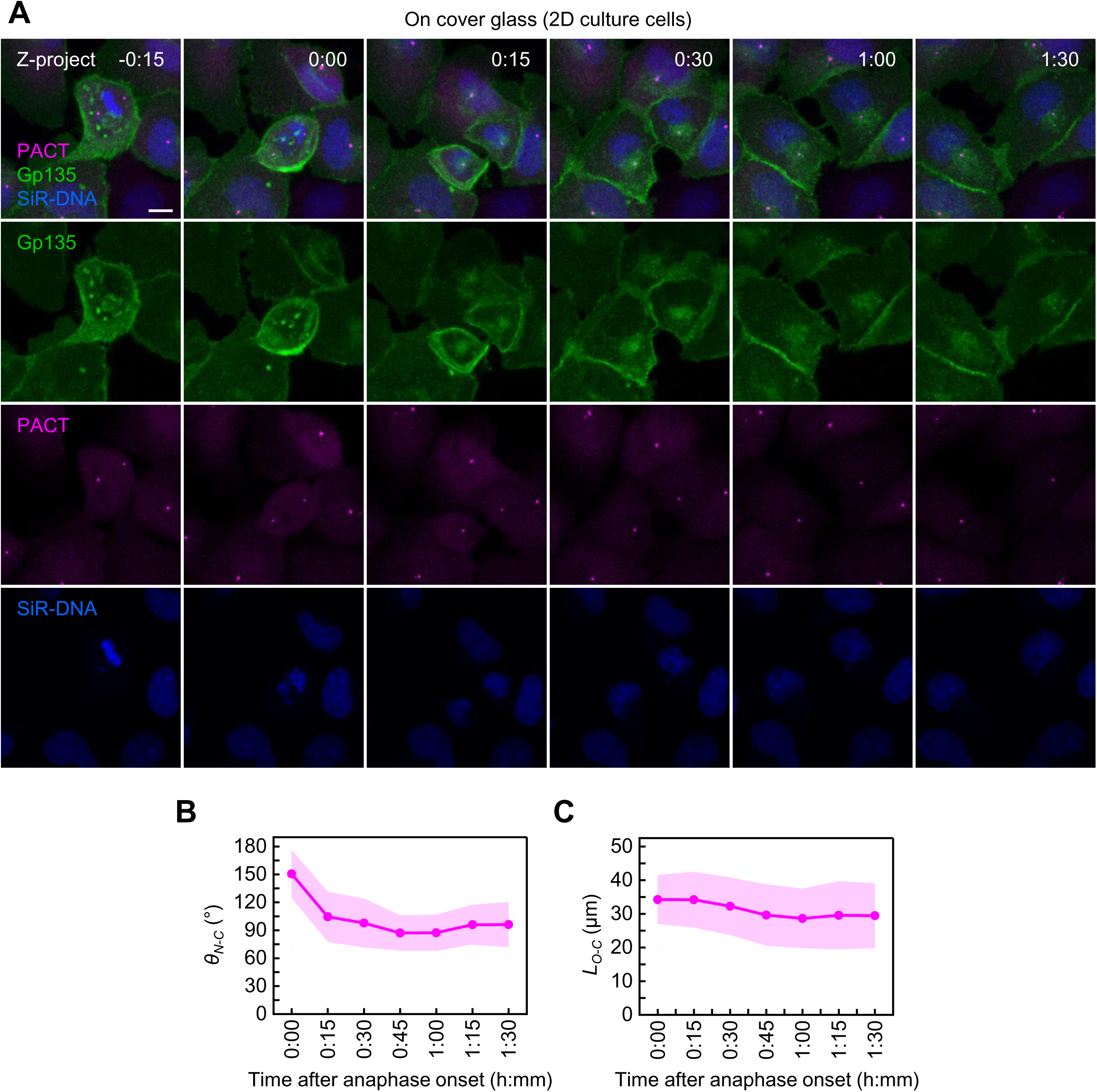
Differential centrosome migration of MDCK cells in adherent 2D culture versus 3D culture conditions during cytokinetic pre-abscission. (A) Time-lapse snapshots of MDCK cells expressing EGFP-Gp135 (green) and PACT-mKO1 (magenta, centrosome marker) on a cover glass. Nuclei were labeled with SiR-DNA (blue) before live imaging. Z-projection images of a dividing cell are shown. Time stamps show hours and minutes, with 0:00 set at the first frame of anaphase onset. Scale bar: 10 μm. (B–C) Change in *θ_N-C_* and *L_O-C_* over time. Each data point represents the average at a given time (6 cells on cover glass from three independent experiments). The lines show the means, and the shaded regions indicate SD values.

**Figure 3—figure supplement 1:**
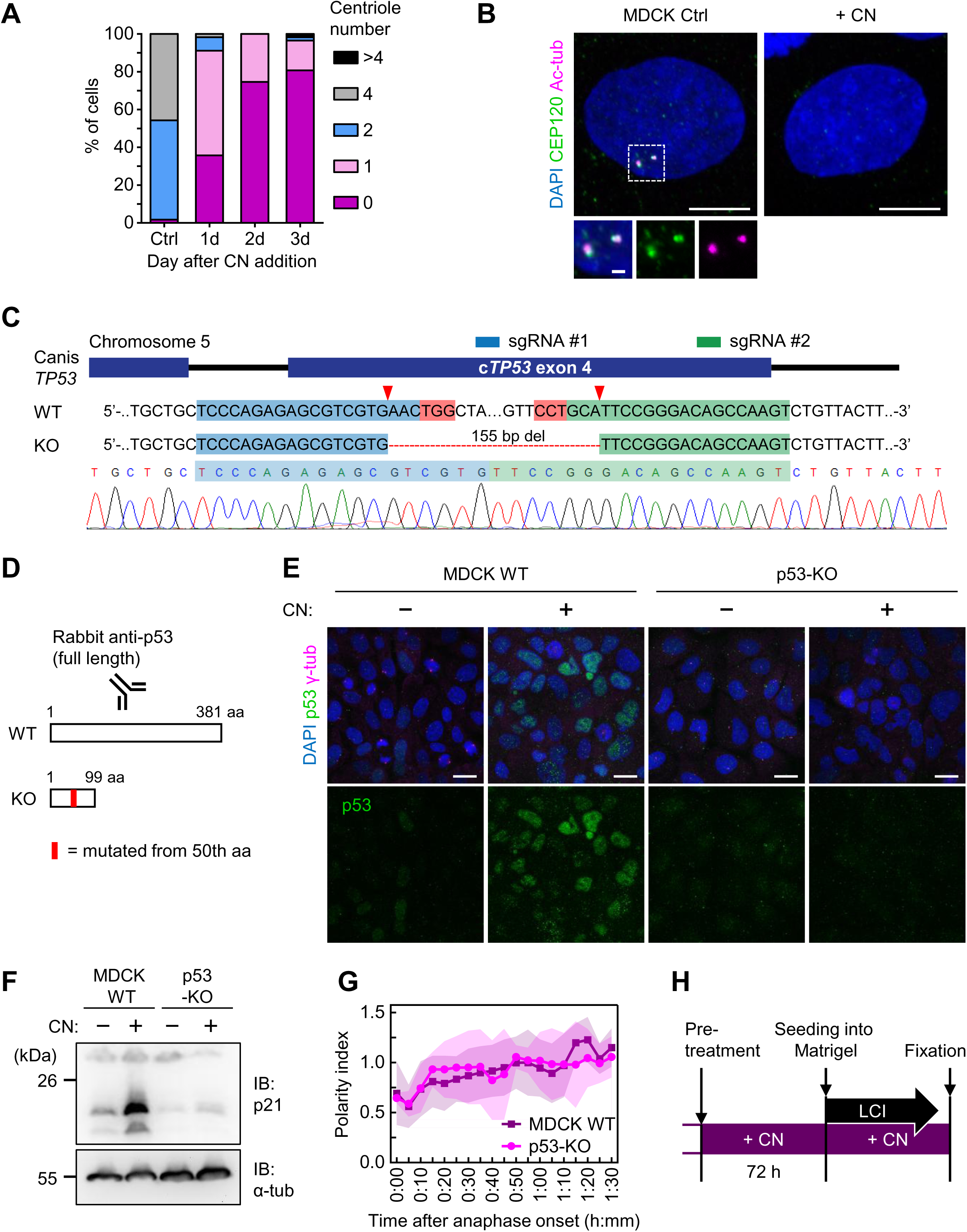
Assessment of centrinone (CN) treatment and generation of p53-KO MDCK cells. (A) Bar graph showing progressive centrosome depletion after CN treatment in MDCK cells. The distribution of centrosome numbers over time after the addition of CN is shown. (B) MDCK cells are present three days after CN addition and an untreated control. Scale bar: 10 μm. Insets show enlargements of the image in the boxed area. Scale bar: 1 μm. (C) Generation of the p53-KO MDCK cell line. The diagram illustrates two single-guide RNA (sgRNA) sequences and their target sites in *p53* exon 4 (blue and green boxes). The red boxes indicate the position of the protospacer-adjacent motif (PAM) sites and the predicted Cas9 cut sites are indicated by red arrowheads. Genotyping results show a genomic mutation in the p53-KO cell line with a deletion (del) within exon 4 of the p53-encoding gene. (D) Binding of p53 antibody and mutation induced in the p53-encoding gene. (E) Immunofluorescence images showing p53 expression in MDCK WT and p53-KO cells with or without CN treatment. Scale bar: 10 μm. (F) Immunoblot results for p21 protein in MDCK WT and p53-KO cells with or without CN treatment. (G) Change in polarity index over time. Each data point represents the average at a given time (ten MDCK WT and four p53-KO cells in Matrigel culture from three independent experiments). The lines show the means, and the shaded regions indicate SD values. (H) Protocol for CN pre-treatment and the time point of Matrigel culture in the presence of CN.

**Figure 3—figure supplement 2:**
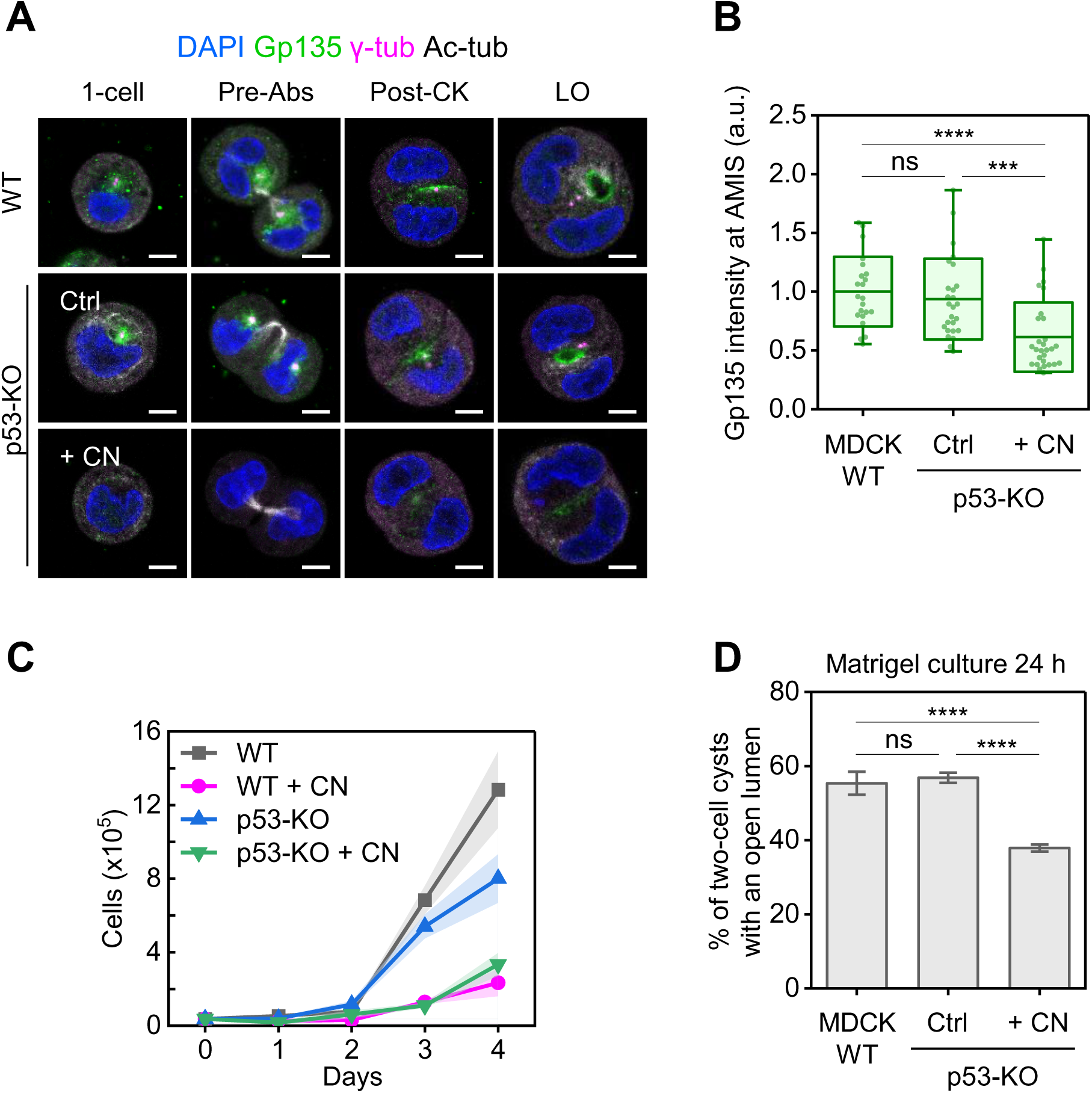
Impact of centriole depletion on MDCK cell proliferation and lumen formation. (A) Single MDCK WT and p53-KO cells, with or without centrinone (CN) treatment, after 8–12 h of Matrigel culture. Single confocal sections through the middle of cysts are shown with immunofluorescent signals of indicated markers: Gp135 (green), γ-tubulin (magenta), acetyl-tubulin (white), and DAPI for nuclei (blue). Acentrosomal cells fail to cluster Gp135-positive vesicles and affect lumen formation in the two-cell stage (Pre-Abs, pre-abscission; LO, lumen open). Scale bar: 5 μm. (B) Boxplot of Gp135 intensity at the AMIS normalized to the mean in WT cells. n = 22 (MDCK WT), 26 (p53-KO), 28 (p53-KO + CN) post-CK cells were analyzed in three independent experiments. Statistical analyses were performed via one-way ANOVA and Dunn’s multiple comparisons (ns: not significant, ***p<0.001, ****p<0.0001). The midlines and boxes show the mean ± SD, with whiskers indicating minimum and maximum values (a.u., arbitrary units). (C) Proliferation curves of MDCK WT and p53-KO cells, with or without CN treatment. The cells with CN treatment underwent a 3-day pre-treatment before examination. Values are presented as the mean ± SD. (D) Proportion of lumen opening in MDCK cell doublets after 24 h of Matrigel culture based on Gp135 staining. n = 17 (MDCK WT), 30 (p53-KO), and 45 (p53-KO + CN) cell doublets were analyzed in three independent experiments. Statistical analyses used two-way ANOVA and Tukey multiple comparisons (ns: not significant, ****p<0.0001). Values represent the mean ± SD.

**Figure 4—figure supplement 1:**
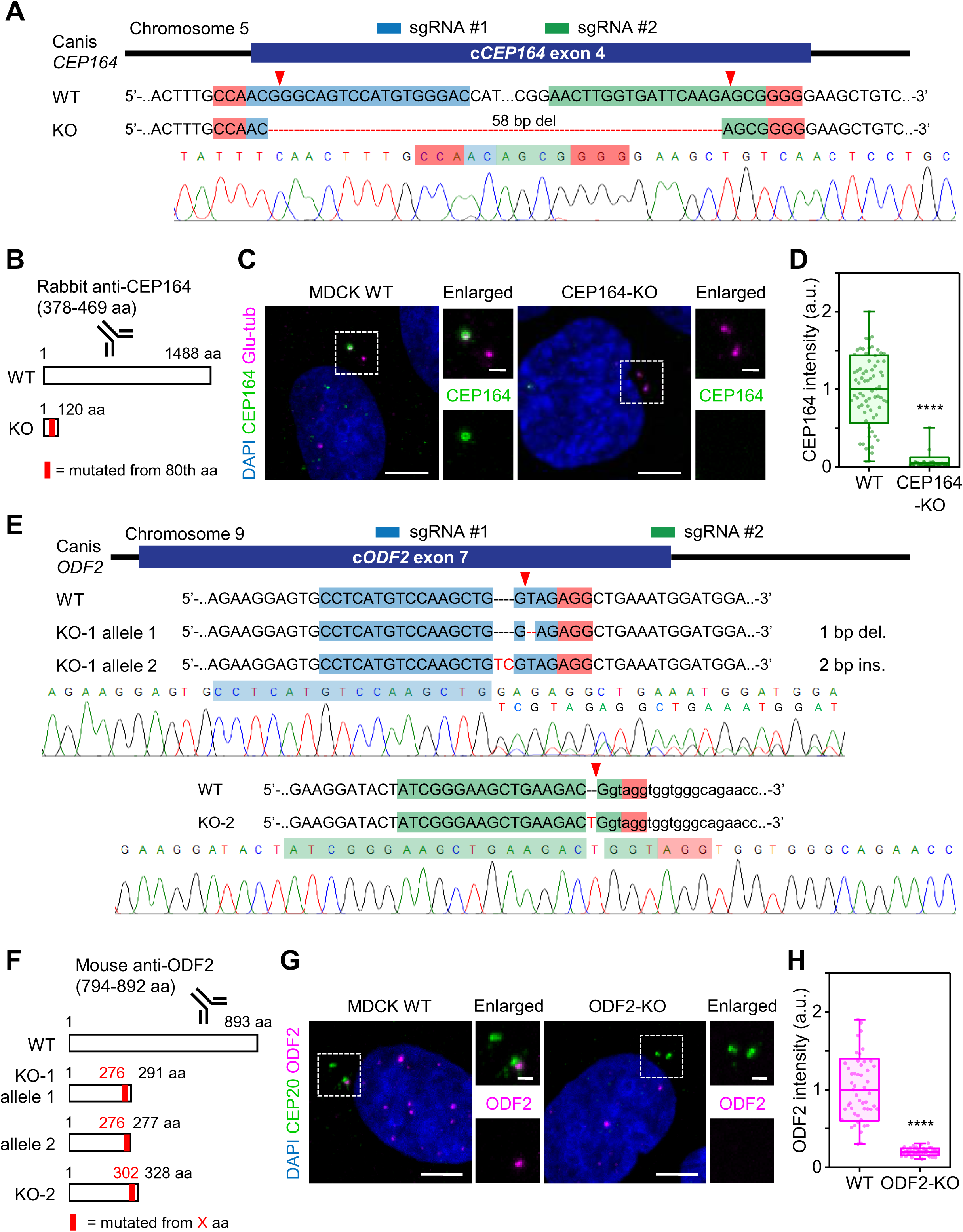
Generation of CEP164 and ODF2-KO MDCK cell lines. (A, E) Diagrams illustrating two single-guide RNA (sgRNA) sequences and their targeting sites within *CEP164* exon 4 and *ODF2* exon 7 (blue and green boxes). The red boxes indicate the position of the protospacer-adjacent motif (PAM) sites and the predicted Cas9 cut sites are indicated by red arrowheads. Genotyping results of the genomic mutations are shown. The predicted genomic mutations (del: deletion, ins: insertion) are illustrated. (B, F) Diagrams illustrating the binding sites of the antibody and the position of the mutation induced in the CEP164- and ODF2-encoding genes. (C, G) Immunofluorescence images depicting CEP164 and ODF2 expression in MDCK WT control and KO cells with glutamylated tubulin and CEP120, respectively, as the centrosome marker. Scale bar: 5 μm. Insets present enlargements of the images within the boxed areas. Scale bar for insets: 1 μm. (D, H) Boxplots of the CEP164 and ODF2 signal at the mother centriole. Fluorescent intensity values of KO cells were normalized to the mean of WT cells. n = 70 (MDCK WT), 51 (CEP164-KO) in (D), and n = 50 (MDCK WT), 50 (ODF2-KO) in (H) were analyzed in three independent experiments. Statistical significance was determined by an unpaired two-tailed Mann–Whitney U test (****p<0.0001). The midlines and boxes show the mean ± SD, with whiskers indicating minimum and maximum values (a.u., arbitrary units).

**Figure 4—figure supplement 2:**
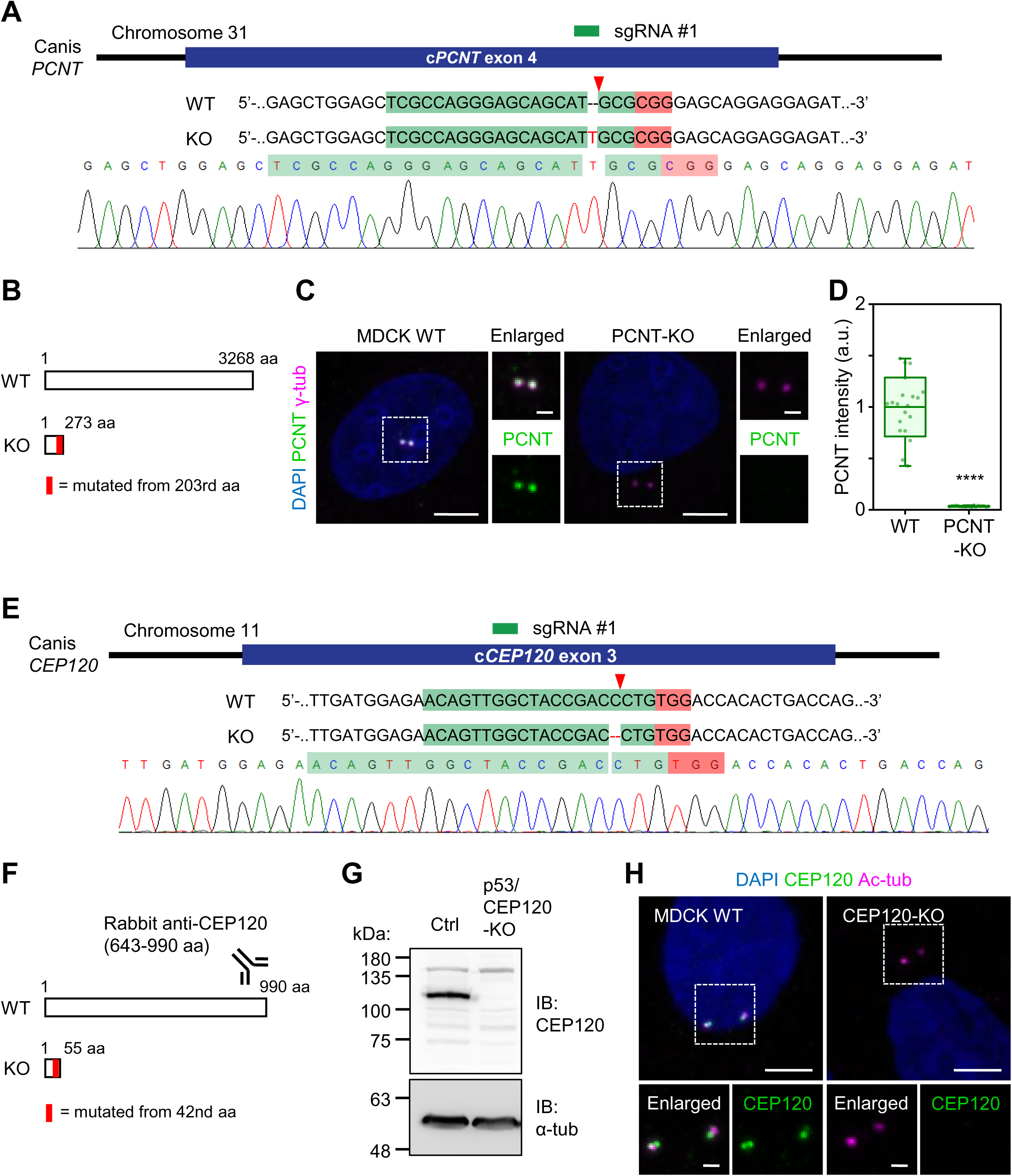
Generation of p53/CEP120 and PCNT-KO MDCK cell lines. (A, E) Diagrams illustrating two single-guide RNA (sgRNA) sequences and their targeting sites within *PCNT* exon 4 and *CEP120* exon 3 (blue and green boxes). The red boxes indicate the position of the protospacer-adjacent motif (PAM) sites, and the predicted Cas9 cut sites are indicated by red arrowheads. Genotyping results of the genomic mutations are shown. The predicted genomi c mutations (del: deletion, ins: insertion) are illustrated. (B, F) Diagrams illustrating the binding sites of the antibody and the position of the mutation induced in the PCNT- and CEP120-encoding genes. (C, H) Immunofluorescence images depicting PCNT and CEP120 expression in MDCK WT control and KO cells with acetyl-tubulin and γ-tubulin, respectively, as the centrosome marker. Scale bar: 5 μm. Insets present enlargements of the images within the boxed areas. Scale bar for insets: 1 μm. (D) Boxplots of the PCNT signal at the centrosome. Fluorescent intensity values of KO cells were normalized to the mean of WT cells. n = 20 (MDCK WT), 20 (PCNT-KO) were analyzed in three independent experiments. Statistical significance was determined by an unpaired two-tailed Mann–Whitney U test (****p<0.0001). The midlines and boxes show the mean ± SD, with whiskers indicating minimum and maximum values (a.u., arbitrary units). (G) Western blot confirming the loss of CEP120 in p53/CEP120-KO cells.

**Figure 4—figure supplement 3:**
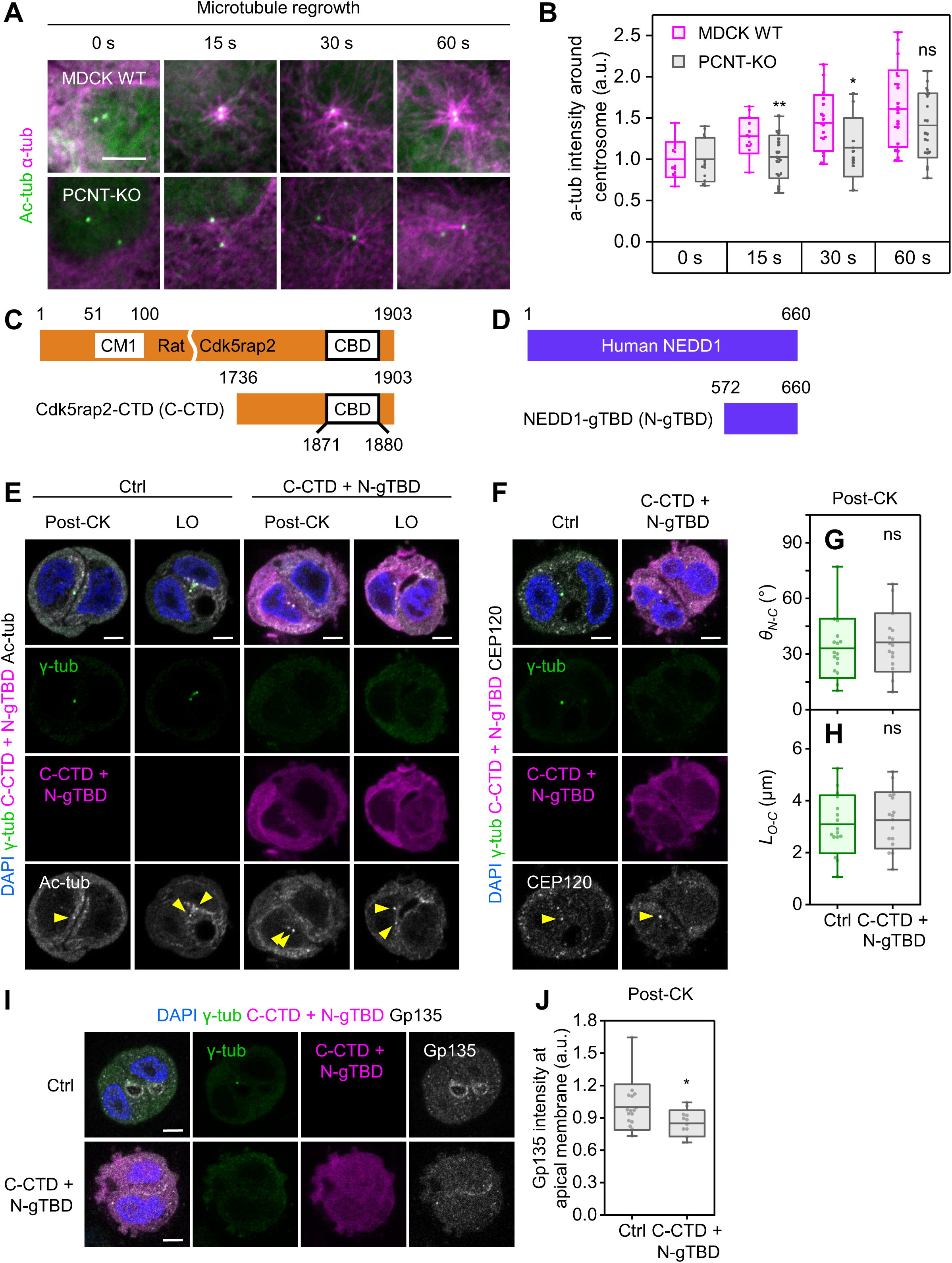
Disruption of centrosomal microtubules impairs epithelial polarization without affecting centrosome positioning in MDCK cells. (A) MT regrowth assays were performed in WT and PCNT-KO cells. Cells were fixed at the indicated times after nocodazole washout and stained for α-tubulin (magenta) and acetylated tubulin (green). (B) Quantification of α-tubulin fluorescence intensity surrounding the centrosome from (A). Fluorescence intensity values at the indicated time points after nocodazole washout were normalized to the mean intensity at 0 s. WT cells: n = 13 (0 s), 13 (15 s), 18 (30 s), and 22 (60 s); PCNT-KO cells: n = 12 (0 s), 20 (15 s), 10 (30 s), and 21 (60 s). Data were pooled from three independent experiments. Statistical significance was assessed using an unpaired two-tailed Mann–Whitney U test (**p<0.01, *p<0.05, ns: not significant). Box plots show the mean ± SD, with whiskers indicating the minimum and maximum values (a.u., arbitrary units). (C) Schematic representation of rat Cdk5rap2 protein and its truncation mutant. CM1 is the γ-TuRC-binding and activating domain. CBD is the centrosome-binding domain. Numbers indicate amino acid (AA) positions. (D) Schematic representation of human NEDD1 protein and its truncation mutant. N-gTBD is the γ-tubulin-binding domain of NEDD1. (E, F, I) Single MDCK cells not expressing (control) or co-expressing mCherry-C-CTD and mCherry-N-gTBD were cultured in Matrigel for 24 h. Images show cells at the post-cytokinesis (CK) and lumen open (LO) stages, with markers including γ-tubulin (green), mCherry (magenta), acetyl-tubulin (white in C), CEP120 (white in D), Gp135 (white in G), and nuclear DAPI (blue). Images are from a single confocal section through the center of the cells. Yellow arrowheads point to the location of the centrosome. Scale bar: 5 μm. (G, H) Boxplots of centrosome positions in control cells (Ctrl; n = 16) and transfected cells (C-CTD + N-gTBD; n = 18) at the post-CK stage, expressed as *θ_N-C_* and *L_O-C_*. Cells were analyzed in three independent experiments. Statistical significance was determined by an unpaired two-tailed Mann–Whitney U test (ns: not significant). Midlines and boxes show the mean ± SD, with whiskers indicating minimum and maximum values. (J) Boxplot of Gp135 intensity on the apical membrane normalized to the mean of control cells. n = 16 (control) and 12 (C-CTD + N-gTBD) cysts were analyzed in three independent experiments. Statistical significance was determined by an unpaired two-tailed Mann–Whitney U test (*p<0.05). The midlines and boxes show the mean ± SD, with whiskers indicating minimum and maximum values (a.u., arbitrary units).

**Figure 5—figure supplement 1:**
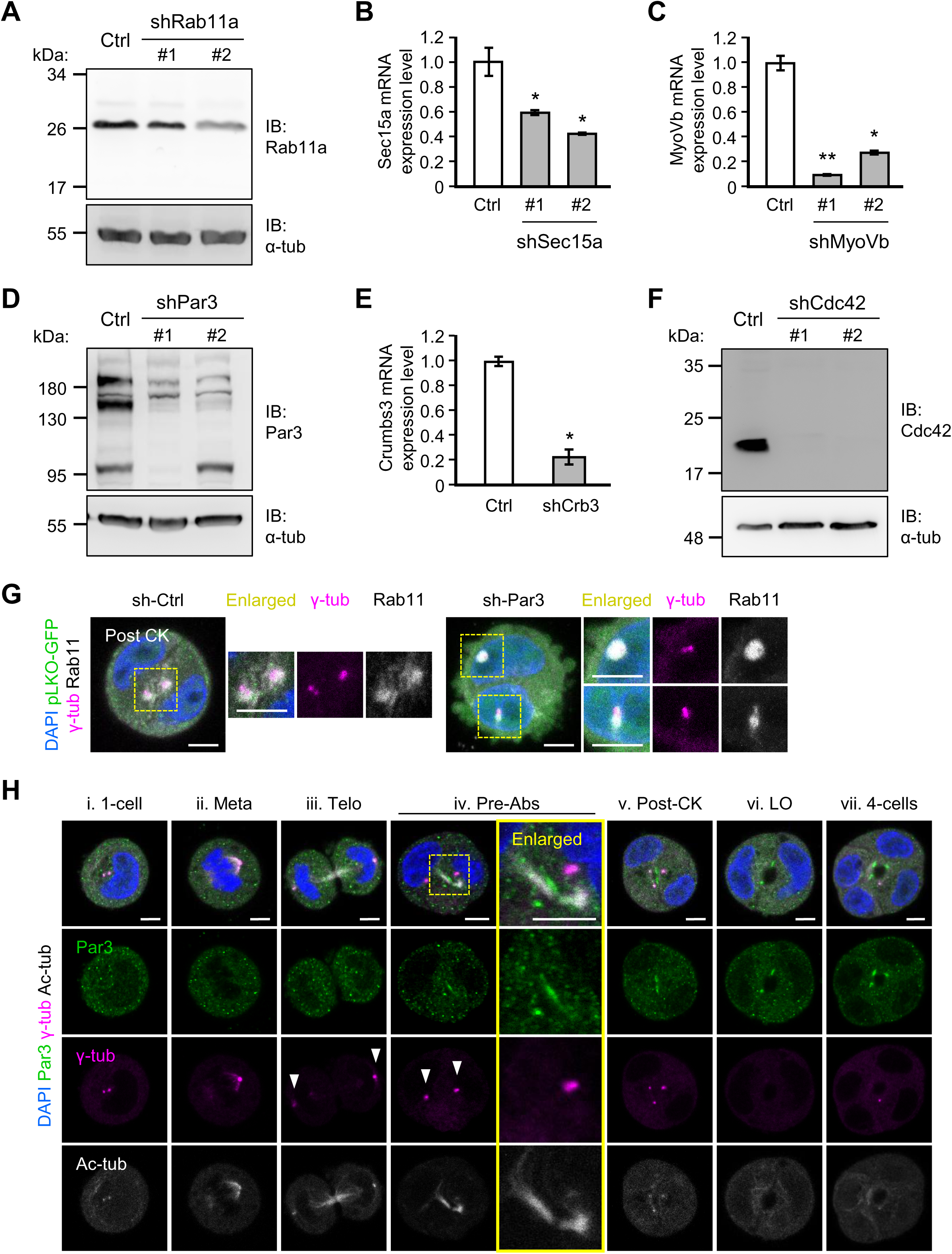
Characterization of MDCK RNAi and localization of Par3 during *de novo* epithelial polarization. (A–F) MDCK cells stably expressing different pLKO-GFP-shRNA constructs targeting Rab11a (A), Sec15a (B), MyoVb (C), Par3 (D), Crb3 (E), and Cdc42 (F). The knockdown efficiency was verified by blotting with appropriate antibodies (A, D, F) or quantitative RT-PCR (B, C, E), with GAPDH used as a control. For quantitative RT-PCR, statistical significance was determined by an unpaired two-tailed Mann–Whitney U test (*p<0.05, **p<0.01). Values represent the mean ± SD. (G) Single MDCK cells stably express pLKO-GFP-shRNA to knock down Par3 after 12 h of Matrigel culture. Images show post-cytokinesis (CK) cells with labeled markers: γ-tubulin (magenta), Rab11a (white), pLKO-GRP (green), and DAPI for nuclei (blue). Z-projection images between two centrosomes are shown. "sh-Ctrl" indicates the expression of scrambled-sequence shRNA. The enlarged images show the centrosome position in the yellow-boxed area. Scale bar: 5 μm. (H) Single MDCK cells after 12 h of culture in Matrigel. Immunostaining signals of the indicated markers are shown: centrosome marker γ-tubulin, polarity protein Par3, acetyl-tubulin, and DAPI. Single confocal sections through the middle of cysts are displayed. The enlarged images display the Par3 signals as they first emerge near the cytokinetic bridge, as indicated in the yellow boxed area. Arrowheads point to the centrosomes (LO, lumen open). Scale bar: 5 μm.

**Figure 5—figure supplement 2:**
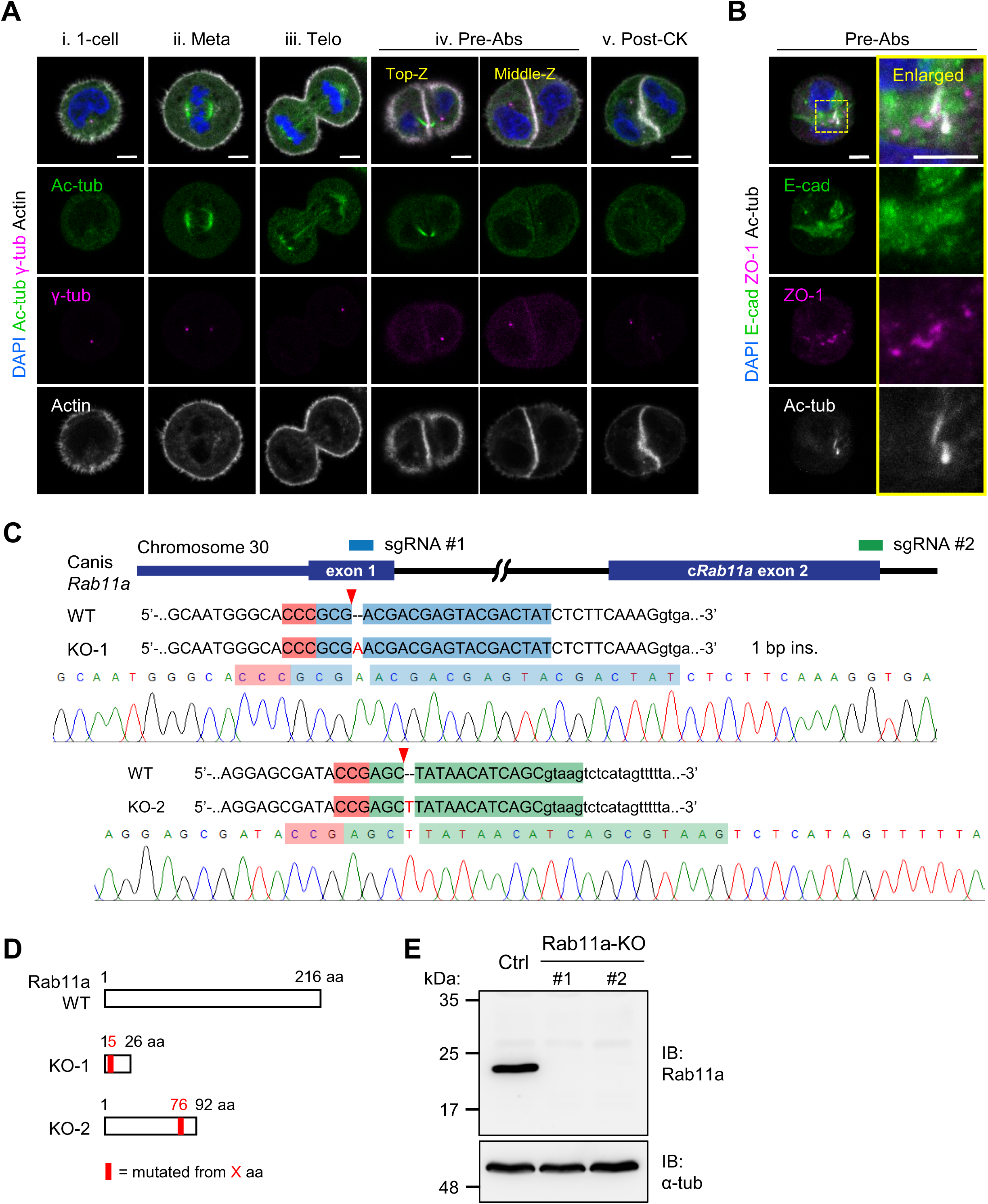
Localization of cell-cell junction components during *de novo* epithelial polarization and generation of Rab11a-KO MDCK cell lines. (A–B) Single MDCK cells after 12 h of culture in Matrigel. Immunostaining signals of the indicated markers are shown: centrosome marker γ-tubulin, polarity protein Par3, acetyl-tubulin, actin, E-cadherin (E-cad), ZO-1, and DAPI. Single confocal sections through the middle of cysts are displayed. The enlarged images display the Par3 and ZO-1 signals as they first emerge near the cytokinetic bridge, as indicated in the yellow boxed area. Arrowheads point to the centrosomes. Scale bar: 5 μm. (C) Diagram illustrating two single-guide RNA (sgRNA) sequences and their targeting sites within *Rab11a* exon 1 and exon 2 (blue and green boxes). The red boxes indicate the position of the protospacer-adjacent motif (PAM) sites and the predicted Cas9 cut sites are indicated by red arrowheads. Genotyping results of the genomic mutations, obtained via gel-purified PCR products covering exon 1 or 2 of the Rab11a-encoding gene in the Rab11a-KO cell line, are shown. The predicted genomic insertion (ins) within exons 1 and 2 of the Rab11a-encoding gene is illustrated. (D) Diagram illustrating the mutation position predicted in the Rab11a-encoding gene. (E) Western blot confirming the loss of Rab11a in two Rab11a-KO lines.

**Figure 6—figure supplement 1:**
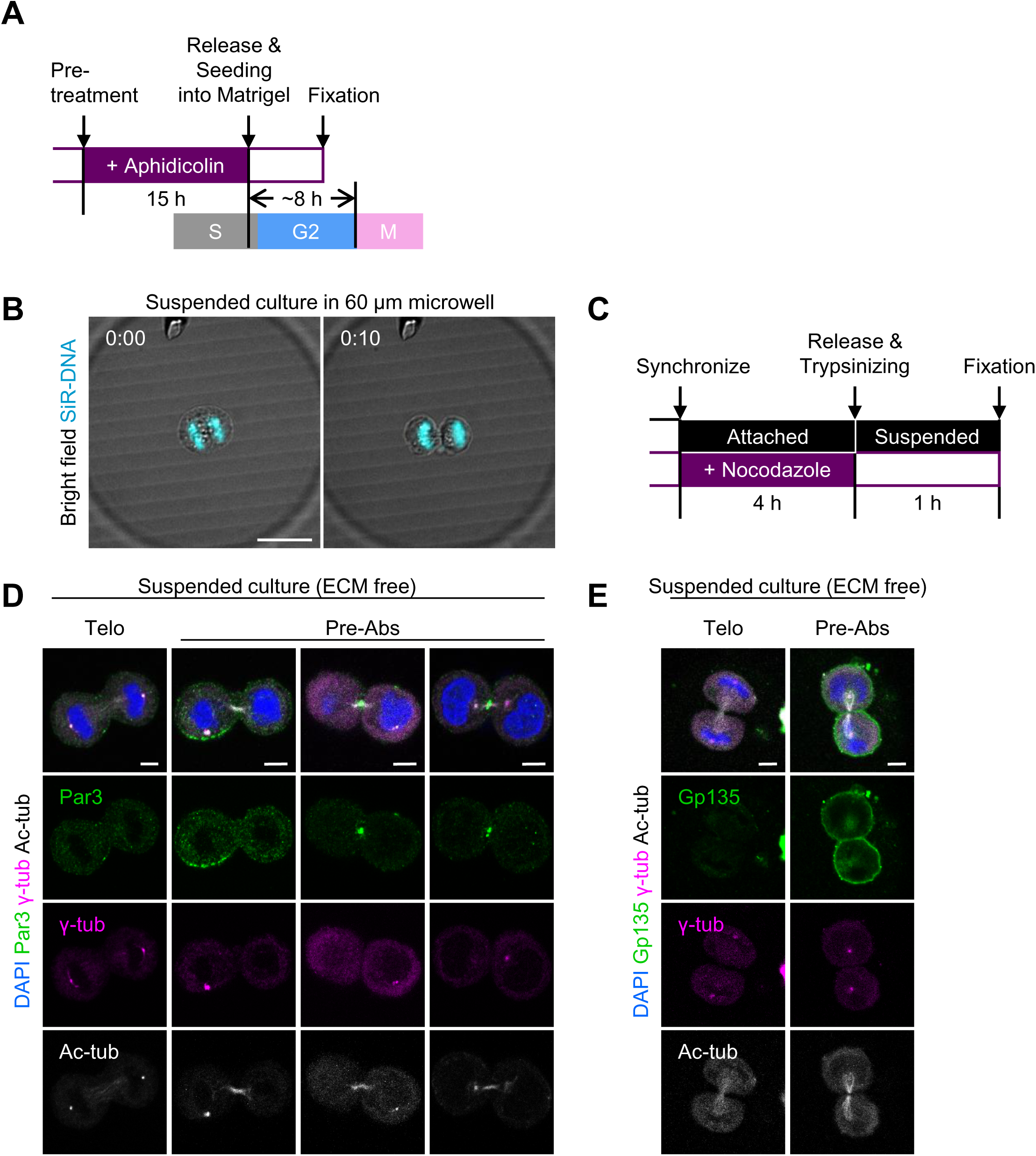
Experimental design and polarity marker dynamics from telophase to pre-abscission in suspension culture. (A) Experimental design: Aphidicolin (Aphi)-treated cells (synchronized at the S phase) were seeded into Matrigel and observed within 8 h to exclude the possibility of cell doublet formation through cell division. (B) Bright-field images of MDCK cells cultured in low-attachment microwells. The images are merged with the DAPI (cyan) signal to show cell division occurring over time. (C) Experimental design: Nocodazole-treated cells (synchronized at prophase) were resuspended in low-attachment dishes and observed within 1 hour to enrich for mitotic cells in suspension culture. (D, E) MDCK cells were treated as indicated in (C), and were labeled with Par3 (green in D), Gp135 (green in E), γ-tubulin (magenta), acetyl-tubulin (white), and DAPI (blue). Images show telophase (Telo) and cytokinetic pre-abscission (Pre-Abs) stages in single a confocal section through the center of a cell doublet.

**Figure 6—figure supplement 2:**
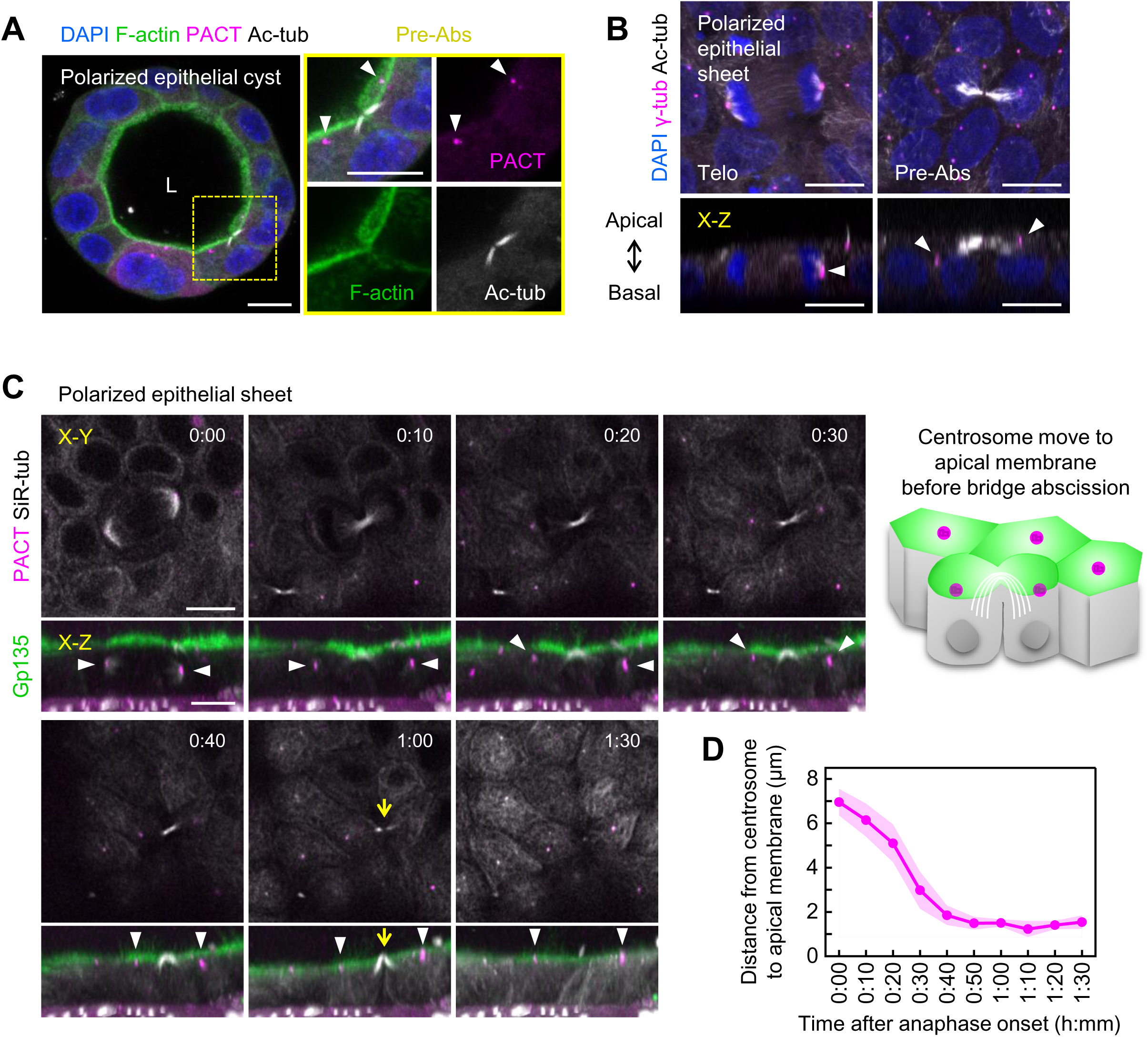
Centrosome position relative to the cytokinesis site in polarized epithelial sheets. (A–B) Polarized MDCK cyst and epithelial sheet with labeled markers: the centrosome markers PACT-mKO1 and γ-tubulin, acetyl-tubulin, F-actin, and DAPI. Insets show enlargements of the image in the boxed area. "L" indicates the lumen in (A). The z-projection of the top view and the X-Z cross-section of the side view (bottom) are shown in (B). Telo, telophase; Pre-Abs, pre-abscission. Arrowheads point to the centrosome position. Scale bar: 10 μm. (C) Time-lapse snapshots of MDCK cells expressing EGFP-Gp135 and PACT-mKO1 cultured on a Transwell insert to form polarized epithelial sheets. The microtubule probe SiR-tubulin (white) was applied before live cell imaging. The X-Y top view and the X-Z cross-section of the side view (bottom) are shown. Time stamps show hours and minutes, with 0:00 set to the first frame after anaphase onset. Arrowheads point to the centrosomes. The yellow arrow indicates the abscission of the cytokinetic bridge. Scale bar: 10 μm. A schematic depicts the centrosomes moving to the apical membrane before bridge abscission. (D) Change in distance from the centrosome to the apical membrane, as measured in each frame (data from three independent experiments, n = 5 cells). The graph presents the mean ± SD.

## Legends to Supplemental Videos

**Figure 2A video: Centrosome migration and apical membrane component trafficking during *de novo* epithelial polarization**

Centrosome migration and apical membrane component trafficking visualized by stable expression of PACT-mKO1 (magenta) and EGFP-Gp135 (green) in Matrigel-cultured MDCK cells. Nuclei were labeled with 500 nM SiR-DNA (blue). The cell was imaged for 1.5 hours (19 frames, 5 min intervals) during the cell division. Time stamps show hours and minutes, with 0:00 set at the first frame of anaphase onset. Scale bar: 10 μm.

**Figure 2—S1A video: Centrosome migration and apical membrane component trafficking in adherent cells**

Centrosome migration and apical membrane component trafficking were visualized by stable expression of PACT-mKO1 (magenta) and EGFP-Gp135 (green) in MDCK cells cultured on a cover glass. Nuclei were labeled with 500 nM SiR-DNA (blue). During cell division, the cell was imaged for ∼4 hours (20 frames, 15-minute intervals). Time stamps show hours and minutes, with 0:00 set at the first frame of anaphase onset. Scale bar: 10 μm.

**Figure 3A video 1: Centrosome migration and apical membrane component trafficking in p53-KO cells**

Centrosome migration and apical membrane component trafficking visualized by stable expression of PACT-mKO1 (magenta) and EGFP-Gp135 (green) in Matrigel-cultured p53-KO cells. Nuclei were labeled with 500 nM SiR-DNA (blue). The cell was imaged for ∼1.5 hours (21 frames, 5 min intervals) during the cell division. Time stamps show hours and minutes, with 0:00 set at the first frame of anaphase onset. Scale bar: 10 μm.

**Figure 3A video 2: Apical membrane component trafficking in centrosome-depleted cells**

Apical membrane component trafficking is visualized by stable EGFP-Gp135 (green) expression in Matrigel-cultured p53-KO cells with centrinone (CN) treatment. Nuclei were labeled with 500 nM SiR-DNA (blue). The cell was imaged for ∼2 hours (24 frames, 5 min intervals) during the cell division. Time stamps show hours and minutes, with 0:00 set at the first frame of anaphase onset. Scale bar: 10 μm.

**Figure 6A video: Centrosome migration and apical membrane component trafficking in aggregated cell doublets**

Centrosome migration and apical membrane component trafficking visualized by stable expression of PACT-mKO1 (magenta) and EGFP-Gp135 (green) in aggregated MDCK cells. Nuclei were labeled with 500 nM SiR-DNA (cyan). The cell was imaged for 3 hours during the contact of two cells. Time stamps show hours and minutes, with 0:00 set at the beginning of live cell imaging. Scale bar: 10 μm.

**Figure 6G video: Centrosome migration and apical membrane component trafficking in suspension-cultured cells**

Centrosome migration and apical membrane component trafficking visualized by stable expression of PACT-mKO1 (magenta) and EGFP-Gp135 (green) in suspension-cultured MDCK cells. Nuclei were labeled with 500 nM SiR-DNA (cyan). Bright-field imaging showed cells suspended in 60 μm diameter microwells. The cell was imaged for ∼1.5 hours during the cell division. Time stamps show hours and minutes, with 0:00 set at the first frame of anaphase onset. Scale bar: 10 μm.

**Figure 6—S2C video: Centrosome migration in polarized epithelial sheets**

Centrosome migration was visualized by stable expression of PACT-mKO1 (magenta) in MDCK epithelial sheets. The apical membrane and microtubule were labeled with EGFP-Gp135 (green) and 750 nM SiR-tubulin (white). During cell division, the cell was imaged for 2 hours (13 frames, 10 min intervals). Time stamps show hours and minutes, with 0:00 set at the first frame of anaphase onset. Scale bar: 10 μm.

**Source data 1.**
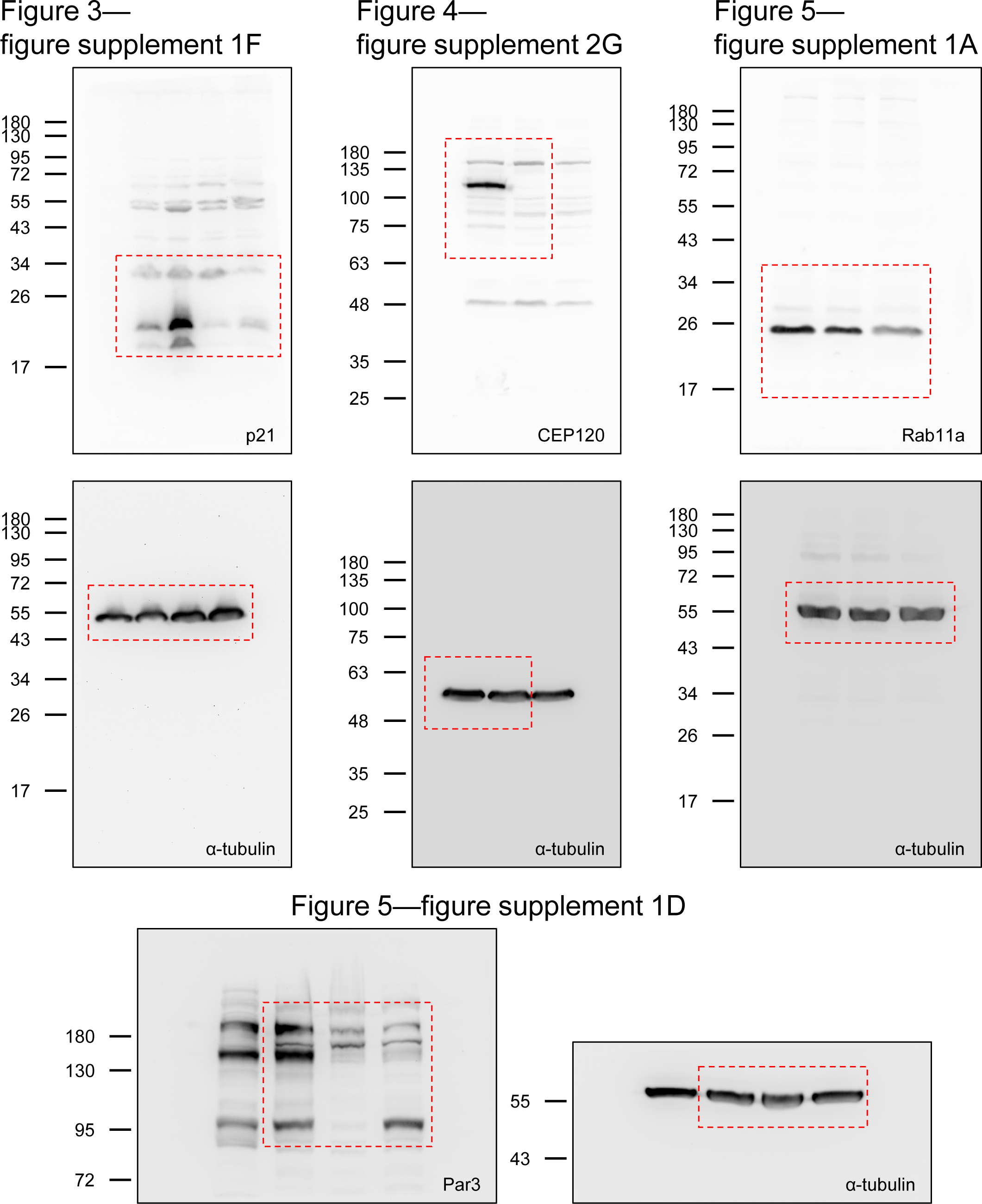
Uncropped Western blots are shown in this manuscript. (A) Western blots showing that p21 is present in MDCK WT and p53-KO cells with or without centrinone (CN) treatment are shown in Figure 3—figure supplement 1F. (B) Western blots showing the knockout of CEP120 from the p53-KO cell line are shown in Figure 4—figure supplement 2G. (C–D) Western blots show the depletion of Rab11a and Par3 in MDCK WT cells, as shown in Figure 5—figure supplement 1A and D.

**Source data 2.**
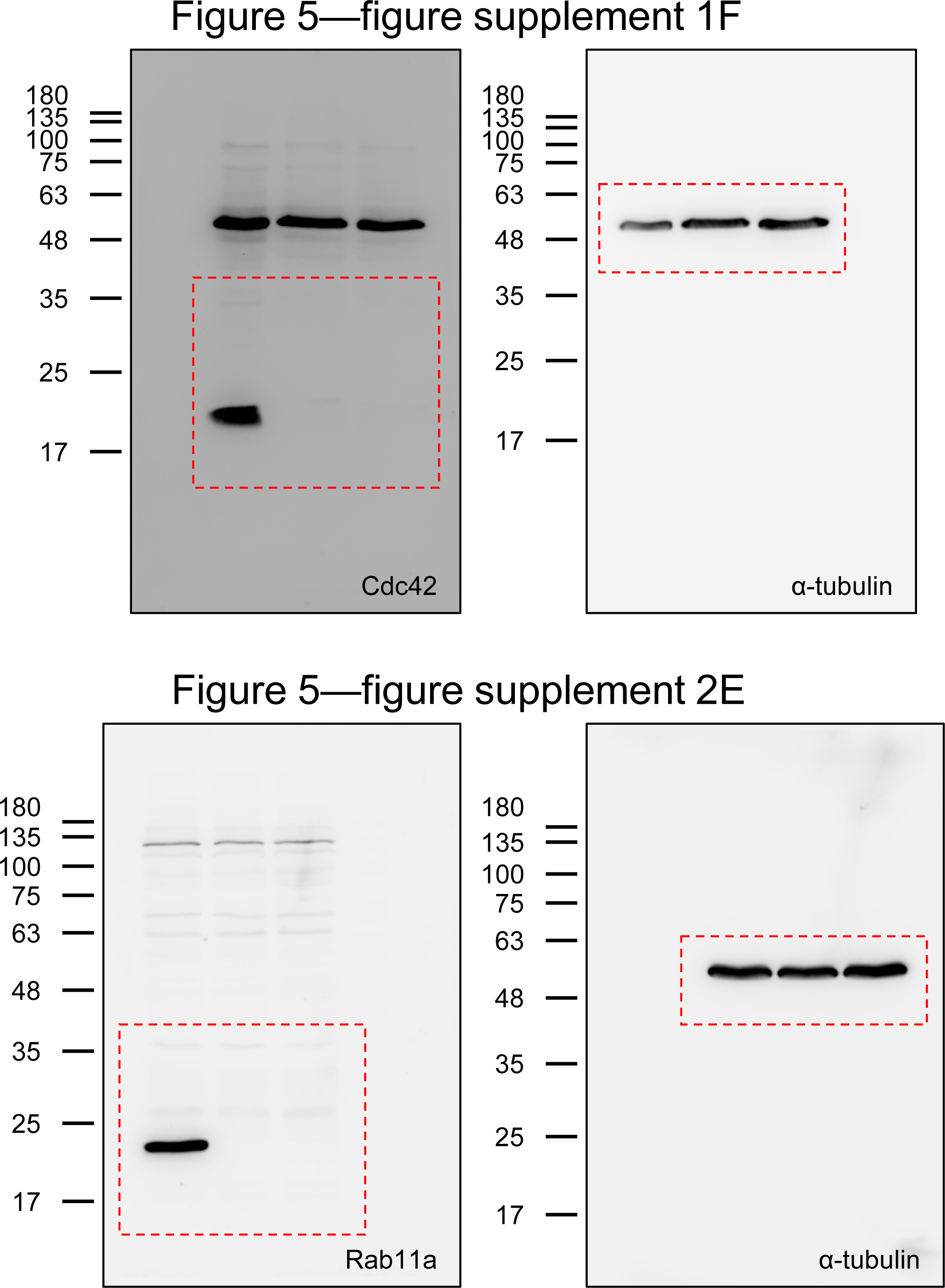
Uncropped Western blots are shown in this manuscript. (A) Western blots showing the depletion of Cdc42 in MDCK WT cells are shown in Figure 5—figure supplement 1F. (B) Western blots showing the knockout of Rab11a from MDCK WT cells are shown in Figure 5—figure supplement 2E.

## Key reagent or resource table

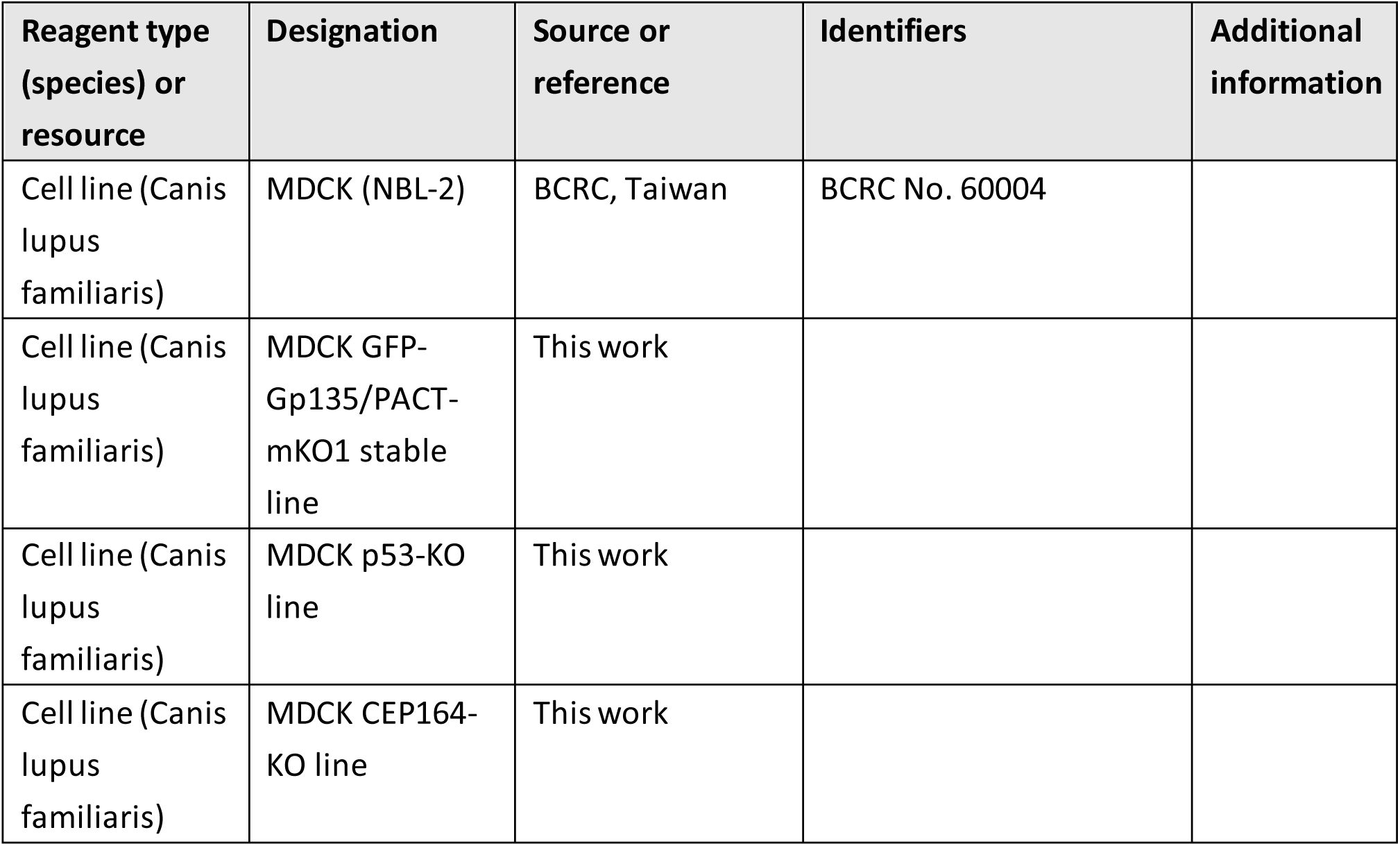
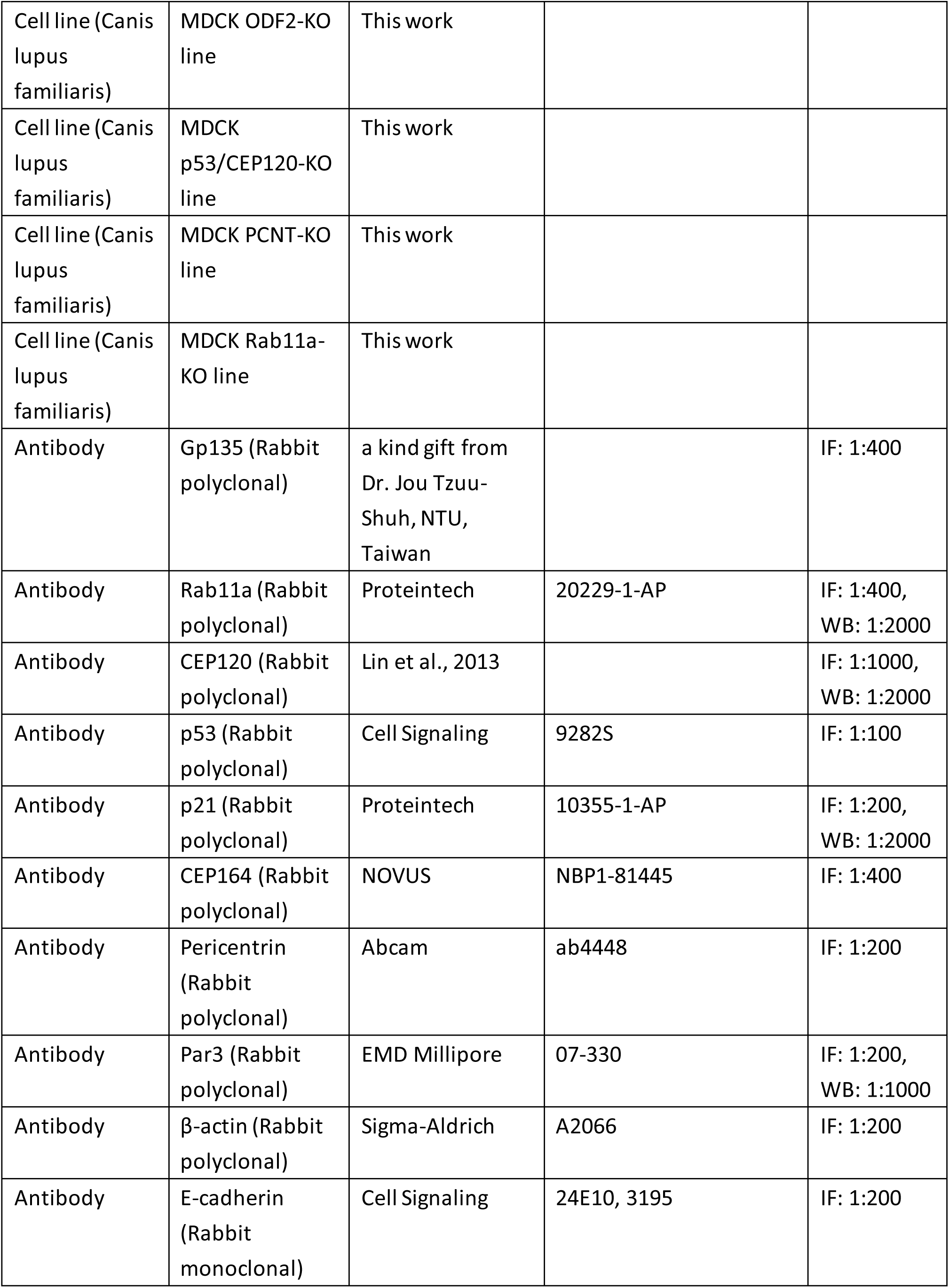
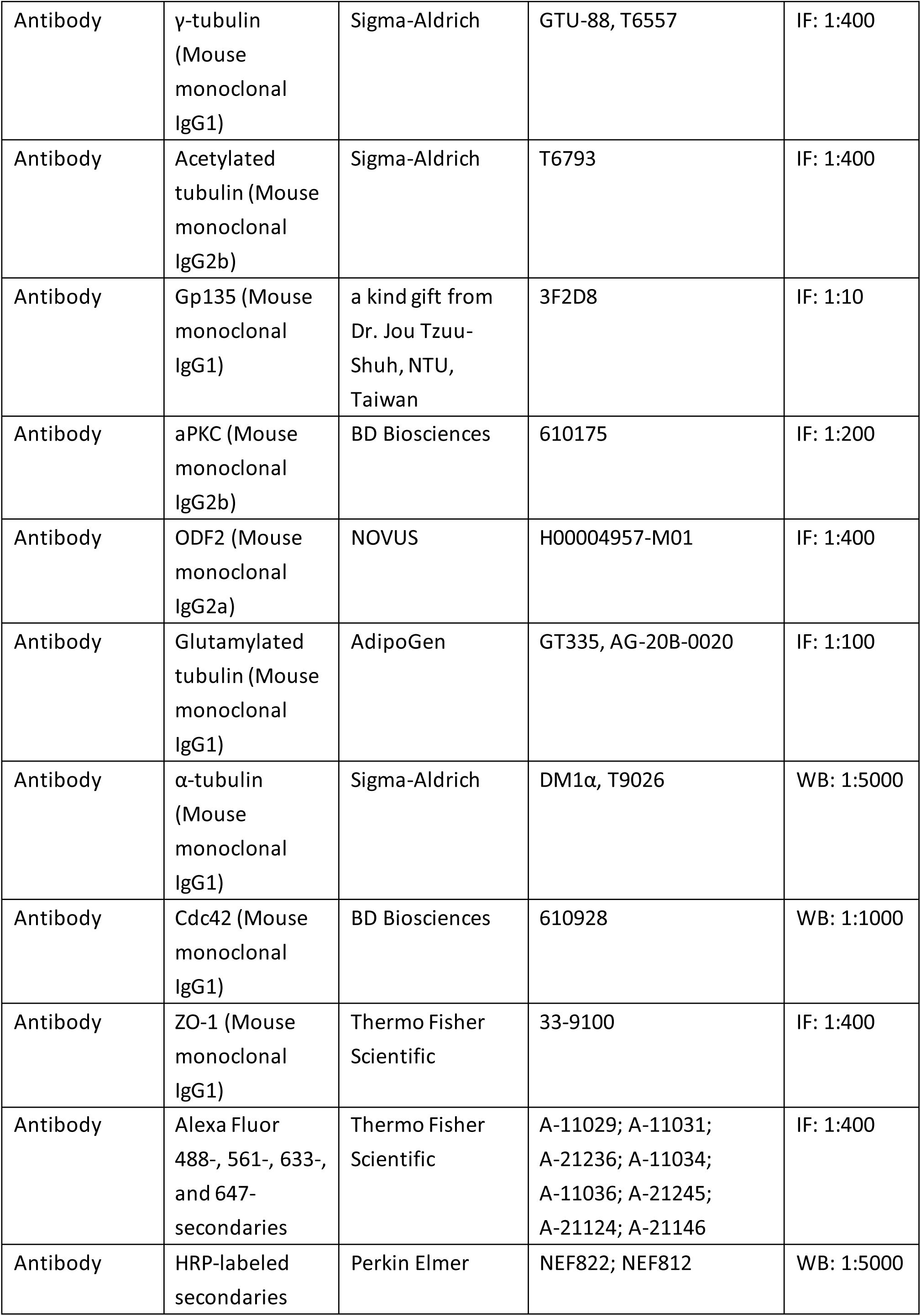
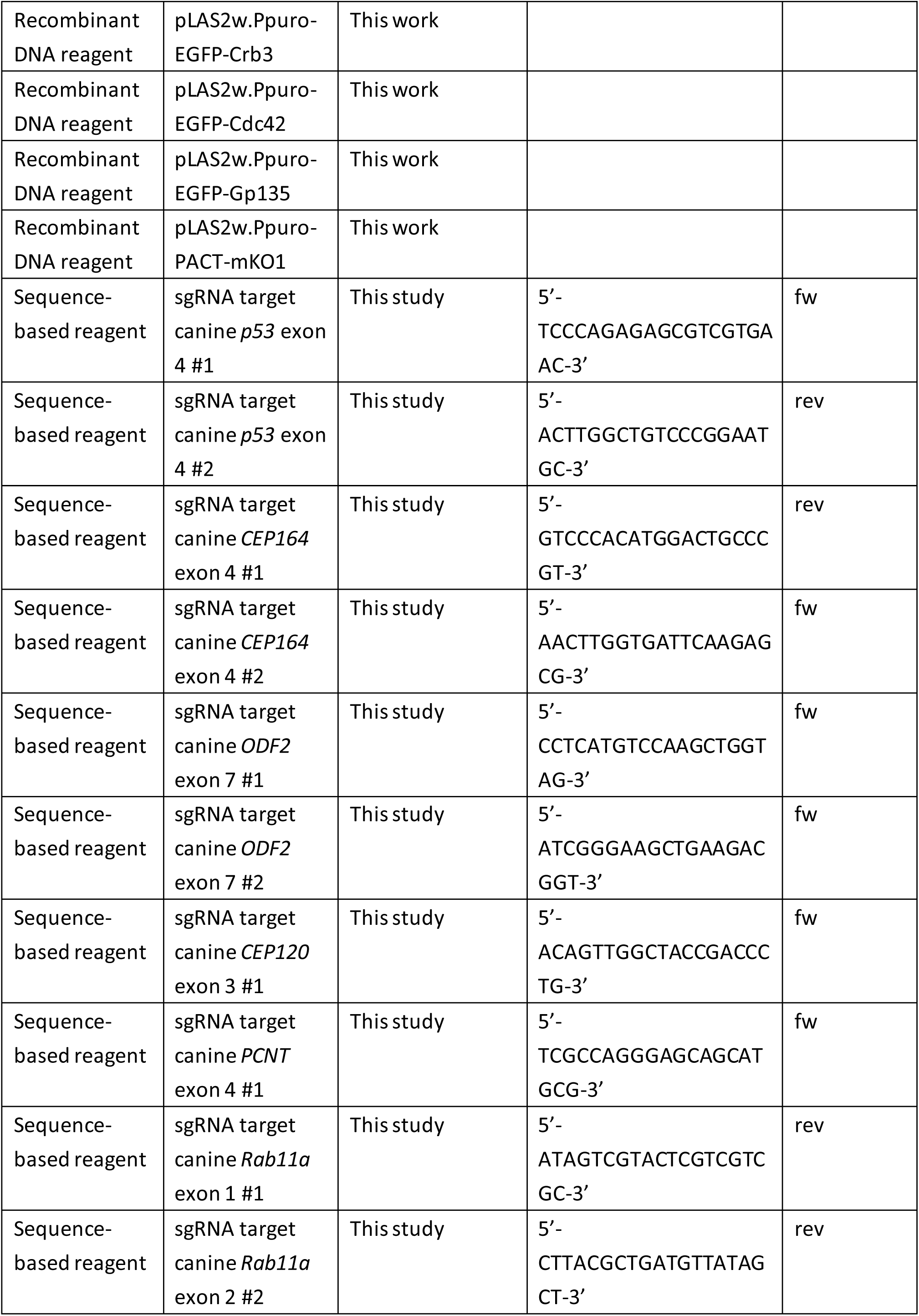
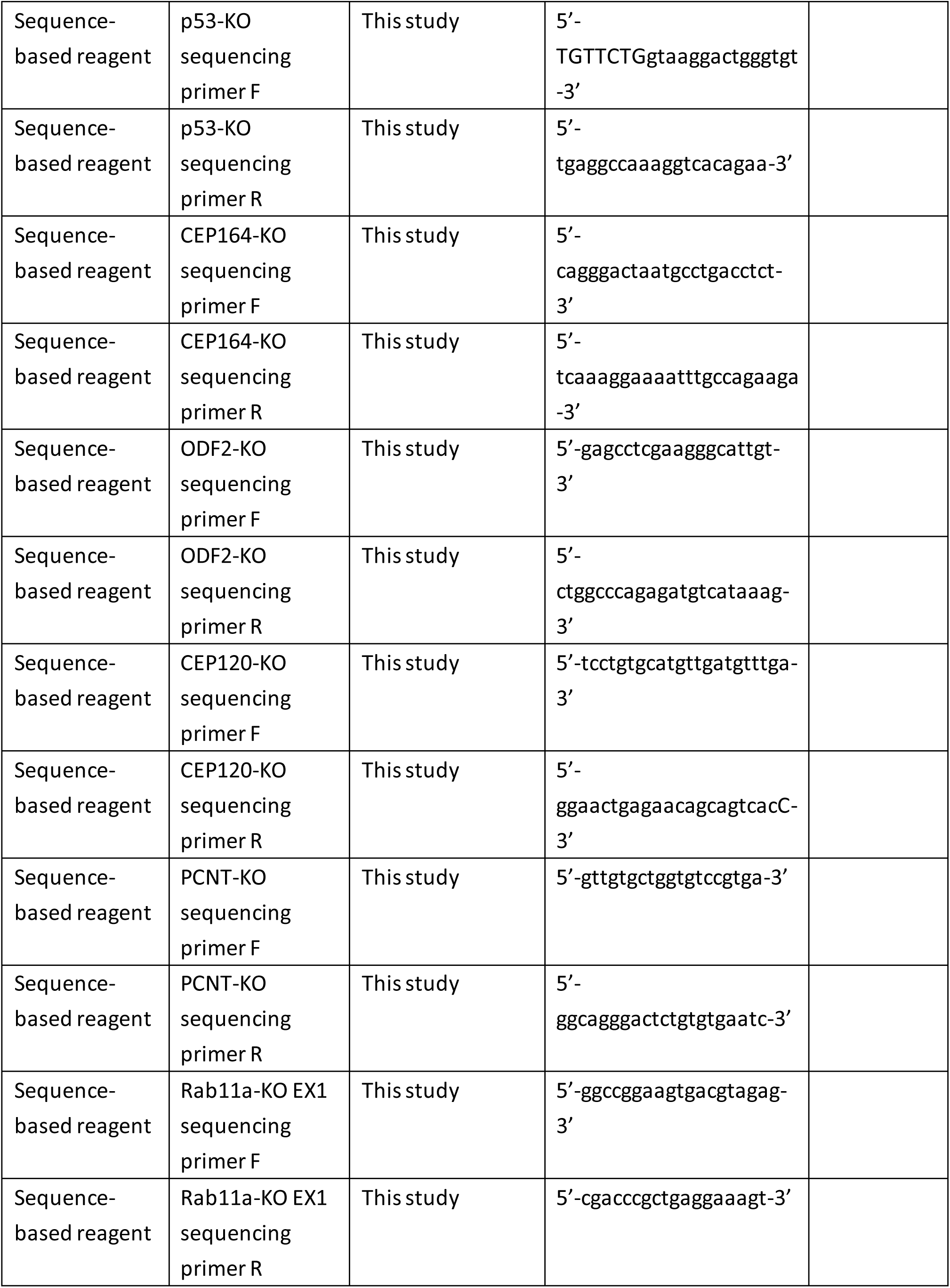
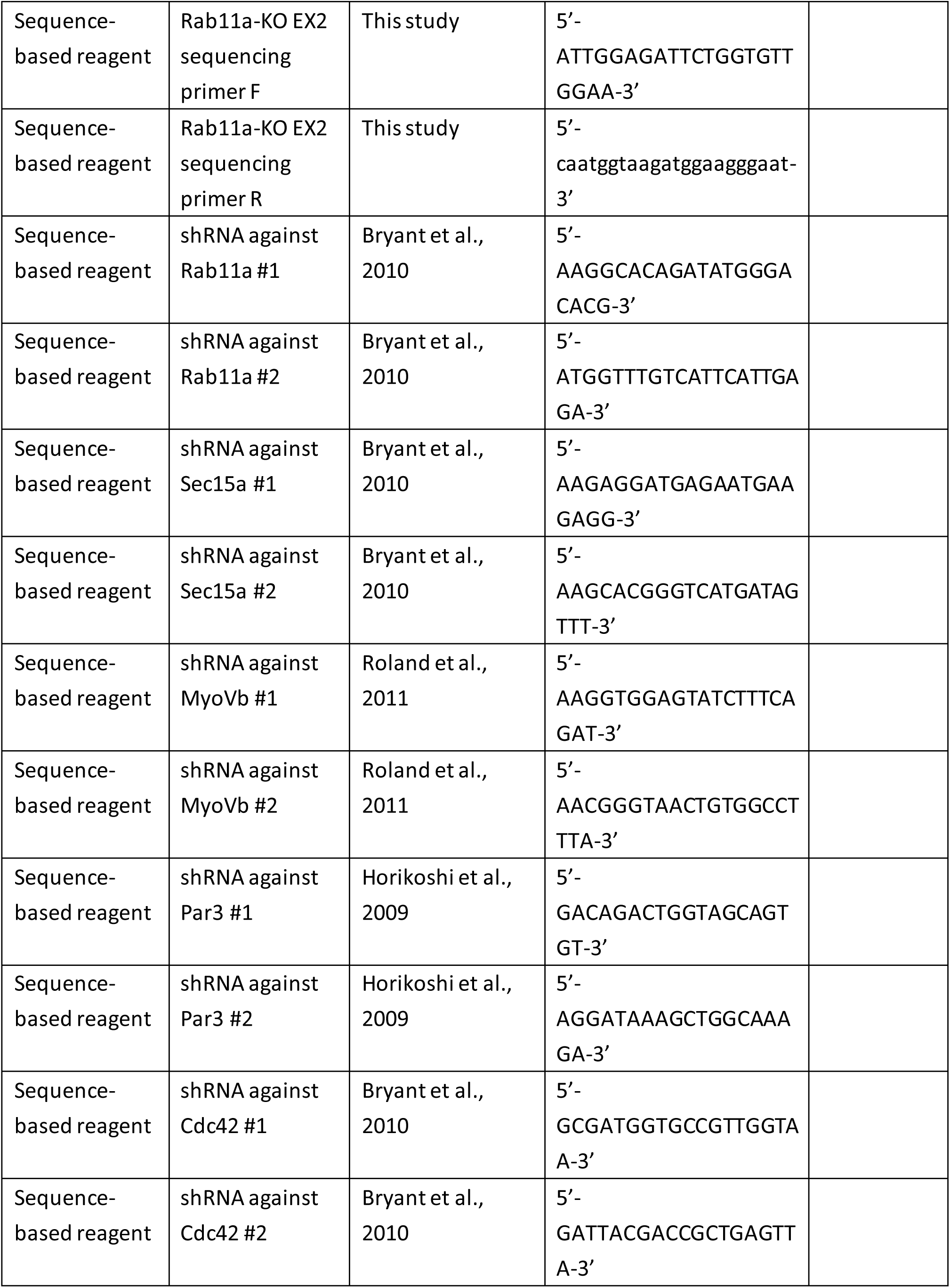
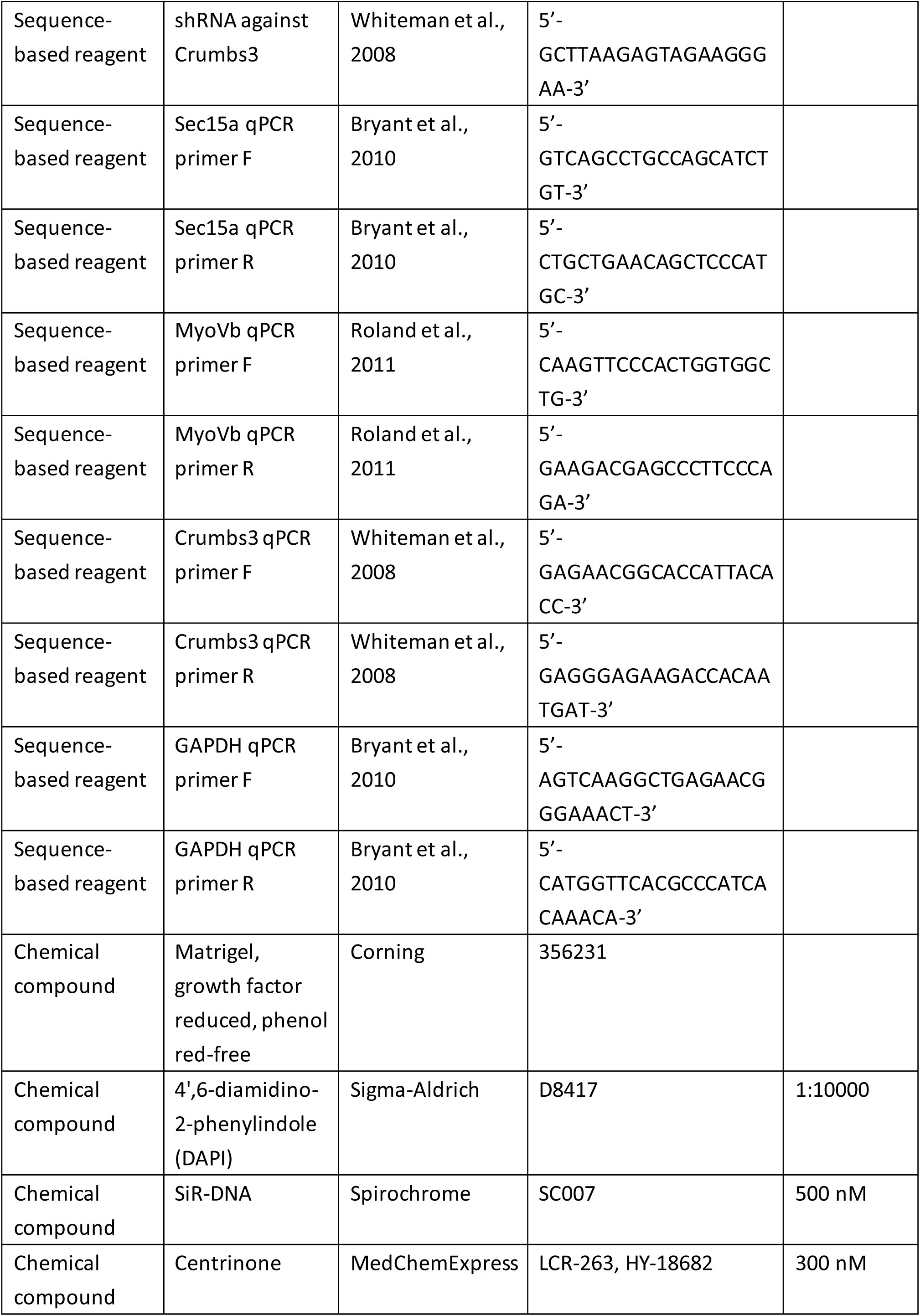
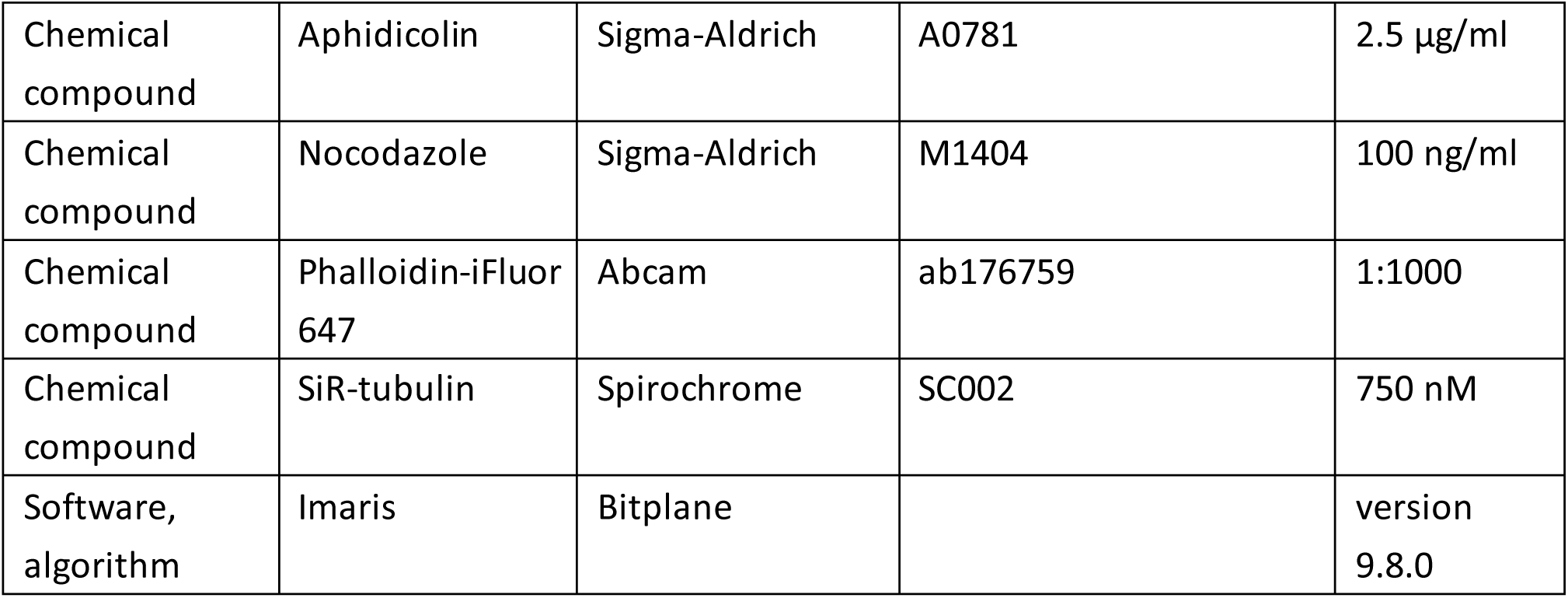

## Notes

### Competing Interest Statement

The authors have declared no competing interest.

### Summary of Updates

We incorporated microtubule regeneration experiments that revealed reduced centrosome microtubule nucleation in PCNT-KO cells (Fig. 4- Supplementary Fig. 3A, B), clarified the interpretation of the polarity index, non-centrosomal GP135 localization, cell-height observations, and ECM-dependent GP135 transport, and corrected figure references where necessary. Furthermore, we expanded the discussion section, comparing our findings with previously described epithelial polarization mechanisms in vivo, including those in the gut of C. elegans.

